# Enhanced CB1 receptor function in GABAergic neurons mediates hyperexcitability and impaired sensory-driven synchrony of cortical circuits in Fragile X Syndrome model mice

**DOI:** 10.1101/2025.01.02.630932

**Authors:** D Gonzalez, CR Jonak, M Bernabucci, G Molinaro, K Collins, SA Assad, JR Gibson, DK Binder, KM Huber

**Affiliations:** Department of Neuroscience, O’Donnell Brain Institute, UT Southwestern Medical Center, Dallas, TX, USA; Division of Biomedical Sciences, School of Medicine, University of California, Riverside, CA, United States of America; Neuroscience Graduate Program, University of California, Riverside, CA, USA

**Author notes:** Contributed equally to this work (D Gonzalez, CR Jonak). co-corresponding authors Electronic address and.

## Abstract

Electroencephalographic (EEG) recordings in individuals with Fragile X Syndrome (FXS) and the mouse model of FXS (*Fmr1* KO) display cortical hyperexcitability at rest, as well as deficits in sensory-driven cortical network synchrony. A form of circuit hyperexcitability is observed in *ex vivo* cortical slices of *Fmr1* KO mice as prolonged persistent activity, or Up, states. It is unknown if the circuit mechanisms that cause prolonged Up states contribute to FXS-relevant EEG phenotypes. Here we examined the role of endocannabinoids (eCB) in prolonged Up states in slices and resting and sensory-driven EEG phenotypes in awake *Fmr1* KO mice. Bidirectional changes in eCB function are reported in the *Fmr1* KO that depend on synapse type (excitatory or inhibitory). We demonstrate that pharmacological or genetic reduction of Cannabinoid Receptor 1 (CB1R) in GABAergic neurons rescues prolonged cortical Up states and deficits in sensory-driven cortical synchrony in *Fmr1* KO mice. In support of these findings, recordings from *Fmr1* KO cortical Layer (L) 2/3 pyramidal neurons revealed enhanced CB1R-mediated suppression of inhibitory synaptic currents. In contrast, genetic reduction of *Cnr1* in glutamatergic neurons did not affect Up state duration, but deletion of *Fmr1* in the same neurons was sufficient to cause long Up states. These findings support a model where loss of *Fmr1* in glutamatergic neurons leads to enhanced CB1R-mediated suppression of GABAergic synaptic transmission, prolonged cortical circuit activation and reduced sensory-driven circuit synchronization. Results suggest that antagonism of CB1Rs as a therapeutic strategy to correct sensory processing deficits in FXS.

## Introduction

Fragile X syndrome (FXS) is the most common form of inherited intellectual disability and a leading monogenetic cause of autism ^1^. Individuals with FXS and the FXS mouse model - the *Fmr1* knockout (KO) display altered sensory processing and sensory hypersensitivity in both the auditory and somatosensory modalities ^2–4^. Electroencephalographic (EEG) recordings in FXS individuals as well as *Fmr1* KO mice provide evidence for enhanced cortical activity at rest, observed as enhanced resting state gamma power^5–9^. Cortical circuits also have impaired temporal processing in response to sensory inputs. This can be observed in FXS individuals and *Fmr1* KO mice as reduced synchronization to time-varying auditory stimuli in the gamma frequency range (40-80Hz), termed intertrial phase coherence or ITPC ^6–9^. Importantly, the severity of EEG alterations in FXS individuals correlates with sensory sensitivities, and deficits in social communication and executive function^7, 10–12^, suggesting the EEG changes are informative clinically. The conserved EEG phenotypes across species suggest a common dysfunction of sensory circuits across FXS individuals and the *Fmr1* KO. Thus, revealing circuit mechanisms of the EEG phenotypes in mice may provide knowledge of circuit dysfunction in humans with FXS.

Findings from acute slices and *in vivo* recordings in *Fmr1* KO mice provide evidence of hyperexcitable cortical circuits that may underlie the EEG phenotypes. Multi-unit recordings *in vivo* in the primary somatosensory or auditory cortex of *Fmr1* KO mice report higher baseline firing rates ^13, 14^. Network hyperexcitability is observed in somatosensory cortex slices or anesthetized *Fmr1* KO mice as an increased duration of persistent activity states, or “Up states”. Up states are a rhythmic oscillation of cortical network activity driven by local excitatory and inhibitory cortical circuits and thus reflect the balanced functioning of excitation and inhibition ^15, 16^. The duration of both spontaneous and thalamically-evoked Up states is prolonged in *Fmr1* KO suggesting that circuit mechanisms that control cortical circuit timing are deficient in the *Fmr1* KO and/or circuits are hyperexcitable ^15,16^. If or how the circuit mechanisms of prolonged Up states contribute to the EEG phenotypes is unclear. The Up state phenotype is robust in a reduced preparation such as the cortical slice and is amenable to probe the cellular and synaptic mechanisms of network dysfunction in *Fmr1* KO.

Pathological endocannabinoid (eCB) signaling through cannabinoid 1 receptors (CB1Rs) has been demonstrated in the *Fmr1* KO and contributes to behavioral phenotypes observed in *Fmr1* KO mice including audiogenic seizures ^17–21^. If or how altered CB1R function contributes to altered sensory cortical networks in the *Fmr1* KO is unknown. A major mechanism by which CB1Rs affect circuit function is through suppression of presynaptic glutamate and GABA release ^22–26^. Activation of postsynaptic Gq-coupled, Group 1 metabotropic glutamate receptors (mGluR1 and mGluR5), ^27, 28^ stimulates synthesis of the eCB, 2-arachidonoylglycerol (2-AG) via diacylglycerol lipase-α (DAGL-α) ^29^. 2-AG traverses the synaptic cleft, activates CB1Rs on presynaptic terminals of glutamatergic and GABAergic neurons which then suppress neurotransmitter release ^22, 23, 30, 31^. Thus, eCBs have the capability to regulate the balance of excitatory and inhibitory synaptic transmission. In the *Fmr1* KO, CB1R-mediated regulation of synaptic transmission is altered, but the direction of change seems to depend on synapse type. In the *Fmr1* KO, mGluR1/5 and CB1R-dependent suppression of inhibitory synaptic transmission is enhanced, whereas suppression of excitatory synaptic transmission is reduced ^17^ ^18–20^. The different CB1R effects on inhibitory vs excitatory synaptic function were performed in different studies and brain regions (hippocampus, striatum, cortex), but effects are consistent for synapse type across regions. Furthermore, both increasing and decreasing eCB/CB1R function have been suggested to be therapeutic strategies for FXS ^17, 21, 32, 33^. The imbalance of CB1R-dependent suppression of inhibitory and excitatory synaptic transmission in the *Fmr1* KO may be expected to lead to hyperexcitability of circuits and results support this view ^18, 21^. However, it is unclear if altered CB1R function in excitatory or inhibitory neurons or both contribute to hyperexcitability related phenotypes associated with FXS.

Towards this goal, we discovered that pharmacological or genetic reduction of mGluR5 activity corrected Up state duration in the *Fmr1* KO ^16^. Because eCB synthesis is a major downstream effector of mGluR5, we investigated the role of eCB synthesis and CB1Rs in prolonged Up states. We hypothesize that the circuit mechanisms of prolonged cortical Up states contribute to the EEG phenotypes of enhanced resting state gamma power and/or deficient ITPC. We test this hypothesis by determining if similar pharmacological and genetic manipulations of CB1R function correct both Up state and EEG phenotypes in *Fmr1* KO mice. Our results indicate that enhanced CB1R-mediated suppression of inhibitory synaptic transmission in *Fmr1* KO cortex contributes to prolonged Up states as well as deficits in cortical circuit synchronization to auditory stimuli *in vivo*. These results provide insight into the cellular and synaptic mechanisms of cortical network hyperexcitability and altered temporal processing. Results suggest that antagonism of CB1Rs may aid sensory processing deficits in FXS.

## Methods and Materials

### Mice

We used the following mouse lines: *Fmr1* KO ^34^, floxed *Fmr1* ^35^, floxed *Cnr1* ^36^, provided by Dr. Joel Elmquist (UT Southwestern Medical Center), VGlut1-Cre (Slc17a7-IRES-Cre; Jackson Labs, stock# 023527) ^37^, and VGlut2-Cre (Slc17a6-IRES-Cre; Jackson Labs, stock# 016963)^38^. All mice were maintained on a C57Bl/6J background by backcrossing with WT mice (Jackson Labs #: 000664). All procedures were approved by the University of Texas Southwestern Medical Center Institutional Animal Care and Use Committee. Experiments were conducted in accordance with the NIH *Guide for the Care and Use of Laboratory Animals*.

### Neocortical Slice preparation, Up state recordings and pharmacological treatments

Up state experiments were performed in acute somatosensory neocortical slices prepared from male (P18-24) littermates of each genotype as described ^16^ (See Supplemental methods). Rimonabant (SR141716A, 5 µM, Tocris, Cat 0923. 0.1% DMSO) and DO34 (10 µM, Glixx Laboratories Inc., 0.1% DMSO) were included in recovery ACSF immediately after slicing and remained in the ACSF during recording (>2 hours).

### Electroencephalogram (EEG)

A multi-electrode (36) array of leads was implanted on the skull surface in P80-90 mice as ^9^ (See Supplemental methods). Resting EEG, sensory responses to Chirp, 40Hz and 80Hz pulses were obtained and analyzed as described ^9^. For rimonabant EEG recordings, approximately 2 hour recordings were performed immediately before and after a 7-day treatment of rimonabant (after is on day 8). Once per day, a dose of 1mg/kg IP was applied as in previous studies ^32, 39, 40^. For the chirp auditory stimulus in these experiments, a 1 second sound intensity ramp preceding the chirp stimulus was not included. The ASSR was the same as for reduced gene dosage experiments except the click train was maintained for 3 seconds.

### RNAscope in situ hybridization (ISH) combined with immunofluorescence

RNAscope was performed in the UT Southwestern Metabolic Core. Mice were anesthetized with a ketamine (80 mg/kg )–xylazine (10 mg/kg) cocktail; i.p.) and transcardially perfused with phosphate buffer saline (PBS) followed by 10% formalin. Brains were post-fixed for 24 hours at 4℃ and then in 30% sucrose for another 24 hours at 4℃. Brains sections (25 µm) were cut with a cryostat and collected in PBS before being treated with hydrogen peroxide for 10 minutes. Sections were mounted onto SuperFrost slides and desiccated overnight at room temperature. On the following day, pretreatment and ISH were performed following the recommendations from the manufacturer (Advanced Cell Diagnostics, USA) and the reagents included in kit cat#323110. The probe for *Cnr1* (cat# 420721-C2) was applied in a solution of probe diluent for 2 hours at 40℃ (HybEZ oven). Slides were incubated with amplification reagents and Opal dyes 570 (1/1,500; Akoya Biosciences). Following the RNAScope procedure, neurons that are GFP-positive were labeled by incubating the brain slides with an antiserum against GFP (Aves Lab; cat# GFP-1020). Slides were next incubated one hour (1/1,000) with a biotinylated anti-chicken secondary (Jackson ImmunoResearch #703-065-155) and one hour (1/1,000) with a streptavidin AlexaFluor488 (Invitrogen, cat# S32354). Slides were rinsed and EcoMount (BioCare Medical, USA) was applied before adding a coverslip.

### Statistical Analysis

Unless stated otherwise, data are presented as the mean ± SEM. Significant differences were determined using *t* tests, one-way ANOVA, two-way ANOVA, when appropriate (all performed with GraphPad Prism). Repeated-measures ANOVA was also used when appropriate. Bonferroni *post hoc* tests were performed following ANOVAs.

## Results

### Antagonism of either CB1R or DAGLα is sufficient to rescue cortical prolonged Up states in Fmr1 KO mice

We examined spontaneously occurring activity states, or Up states, in acutely prepared brain slices containing somatosensory cortex. Up state duration is longer in *Fmr1* KO mice, and we interpret this to reflect circuit hyperexcitability in the cortex ^15^. To determine if longer Up states in *Fmr1* KO mice are mediated by altered eCB signaling, we pharmacologically blocked eCB signaling. If longer Up states involved enhanced eCB signaling – like that occurring at GABAergic synapses – then blocking CB1Rs would be expected to reduce Up state duration to normal WT levels. We refer to this as a “rescue.” On the other hand, if the long Up states were a consequence of diminished eCB signaling – like that reported for glutamatergic synapses – then blocking eCB signaling may either prolong Up states or have no effect in the *Fmr1* KO.

As a first test, we blocked CB1Rs by preincubating slices (2 hrs) with the CBR1 an inverse agonist, rimonabant (5 µM, 0.1% DMSO), or vehicle (0.1% DMSO). In vehicle treated slices, Up states were longer in *Fmr1* KO slices compared to WT, as expected (Fig. 1A,B,C). Rimonabant treatment reduced Up state duration in the *Fmr1* KO as compared to vehicle-treated controls (Fig. 1B,C). This result is consistent with enhanced, eCB-mediated suppression of GABAergic transmission contributing to long Up states in the *Fmr1* KO. Interestingly, rimonabant had no effect on Up state duration or frequency in WT slices, but reduced Up state amplitude across both genotypes (2 way ANOVA, Main effect of rimonabant; F (1, 109) = 4.071; p< 0.05; Fig. Supplementary (S) 1) which was similar to previous observations ^41^.

**Figure 1.**
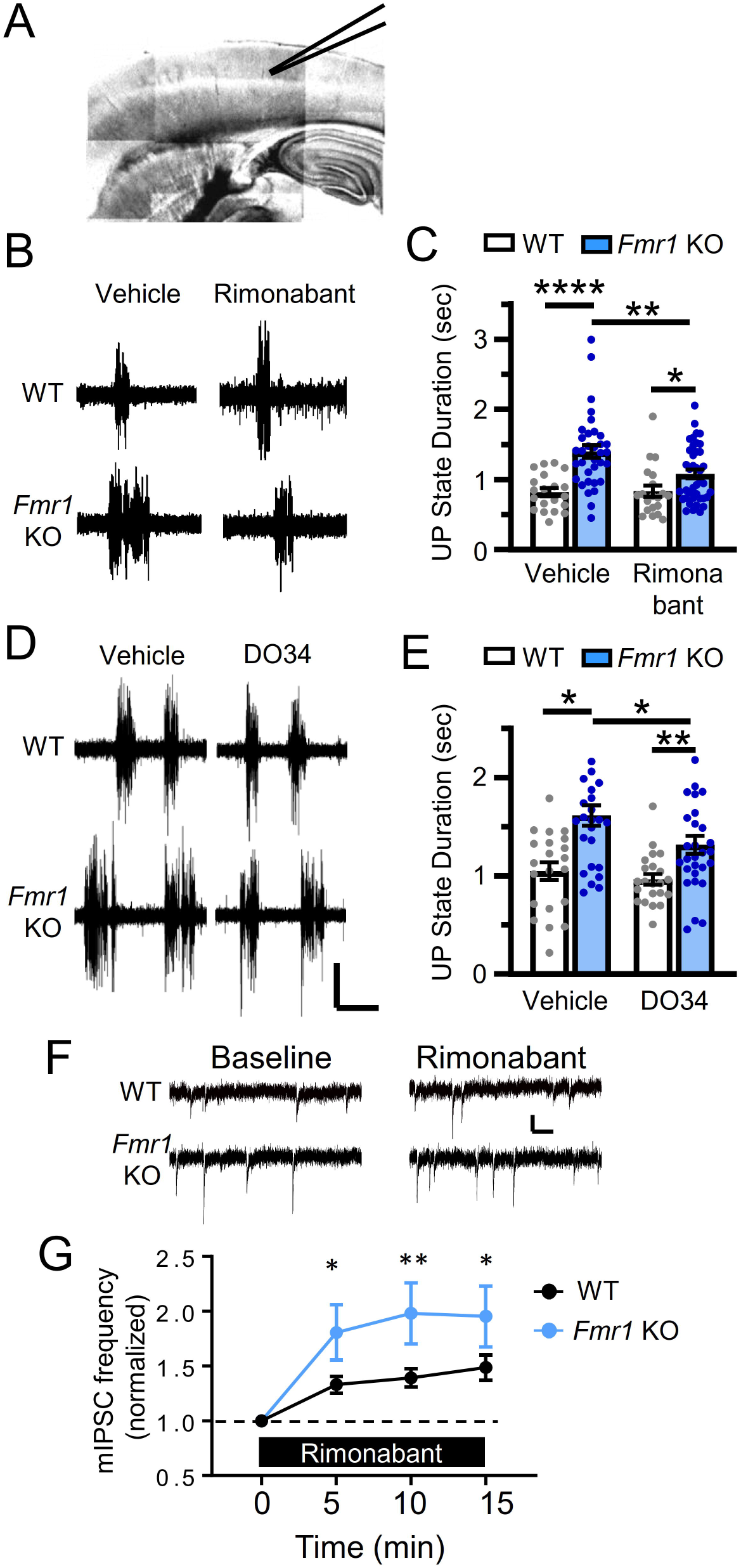
Antagonism of CB1Rs and DAGLα rescues prolonged Up states in *Fmr1* KO cortical slices. A) Up states were recorded extracellularly in layer 4 of somatosensory cortex. B) Example traces collected in vehicle and the CBR1 antagonist, Rimonabant (5 µM). C) Bar-graph of Up state duration showing a prolonged phenotype in vehicle and no detectable change in the *Fmr1* KO with Rimonabant application (N=20-42 slices, 5-9 mice for each condition). D) Example traces collected in vehicle and the diacylglycerol lipase inhibitor, DO34 (10 µM). E) Bar-graph of Up state duration showing a prolonged phenotype in vehicle and a partial rescue in the *Fmr1* KO with DO34 application (N=21-29 slices and 5-7 mice for each condition). Scale bars for B and D = 50 µV and 1 s. 2-way ANOVA with Sidak’s multiple comparisons test. F,G) Enhanced CB1R-mediated suppression of inhibitory synaptic transmission in *Fmr1* KO cortex. F) Examples mIPSCs from a L2/3 pyramidal neuron before (baseline) and after Rimonabant (5 µM) application. Scale bars = 5 pA and 50 ms. B) Average mIPSC frequency (normalized to pre-rimonabant baseline) in WT and *Fmr1* KO neurons. (N=14-20 cells from 7-15 mice per genotype). Repeated measures 2 way ANOVA; Rimonabant X genotype interaction (F (3, 96) = 3.366; p<0.05); Sidak’s posthoc tests; *, p<0.05 **, p<0.01 ****, p<0.001.

mGluR5 stimulates synthesis of the eCB 2-Arachidonoylglycerol (2-AG) by generating its precursor diacylglycerol. Diacylglycerol is then converted to 2-AG by diacylglycerol lipase-α (DAGLα) ^42^. To test if 2-AG synthesis was required for long Up states, we pretreated slices with a selective DAGLα inhibitor, DO34 (10 µM, 0.2% DMSO), or vehicle (0.2% DMSO) ^43–45^. As observed for rimonabant, DO34 treatment reduced Up state duration in the *Fmr1* KO (Fig. 1D,E) and had no effect on duration in WT slices. DO34 also did not affect Up state frequency or amplitude in either genotype (Fig. S1C,D). These results are consistent with enhanced, ongoing eCB-mediated suppression at GABAergic synapses in the *Fmr1* KO that leads to longer Up states.

### Changes in GABAergic synaptic currents with application of a CB1R antagonist are consistent with greater eCB-mediated suppression in the Fmr1 KO

The rescue of Up state duration with rimonabant suggests that CB1Rs are driving circuit hyperexcitability. We hypothesized that the CB1R-mediated suppression of inhibitory synaptic transmission may be enhanced in *Fmr1* KO cortex similar to what has been observed in striatum and hippocampal CA1 ^18, 19^. We tested the hypothesis by examining the effects of rimonabant on spontaneous miniature (m) IPSCs. While CB1Rs typically suppress evoked synaptic transmission ^46^, effects can also be observed on mIPSCs in the presence of Ca^2+^ ^47–49^. We measured mIPSC amplitude and frequency before and after wash-in of rimonabant (5 µM, Fig. 1F,G, 0.1% DMSO). Baseline mIPSC frequency and amplitude was not different between WT and *Fmr1* KO mice (Fig. S1E,F). If chronic, CB1R-mediated suppression of GABAergic transmission exists in the *Fmr1* KO, a greater increase in either mIPSC amplitude or frequency would be expected in *Fmr1* KO slices with rimonabant. We observed an increase in mIPSC frequency in both WT and *Fmr1* KO slices, but the effects of rimonabant on mIPSC frequency were greater in the *Fmr1* KO (Fig. 1G; 2way ANOVA; Main effect of genotype; F (1, 32) = 4.598; p<0.05; genotype X time interaction; F (3, 96) = 3.366; p< 0.05). This result is consistent with enhanced CB1R-mediated suppression of inhibitory synaptic transmission in the *Fmr1* KO cortex and suggests that this mechanism contributes to network hyperexcitability.

### Genetic reduction of Cnr1 in GABAergic neurons, but not glutamatergic neurons, rescues prolonged Up states in Fmr1 KO mice

To determine if enhanced function of CB1Rs in inhibitory neurons contributes to prolonged Up states in *Fmr1* KO, we conditionally deleted *Cnr1* in GABAergic neurons using the VGAT-Cre mouse line. To confirm this genetic targeting strategy, we bred *Cnr1* floxed mice (*Cnr1^fl/fl^*) with both *VGAT*^Cre^ and Rosa26-EYFP Cre reporter (*Rosa26*^EYFP^) mice and performed RNAscope for *Cnr1* together with immunohistochemistry for EYFP to mark Cre expressing GABAergic neurons. On the wildtype background, (*VGAT*^Cre^:*Rosa26*^EYFP^:*Cnr1*^+/+^*) Cnr1* expression is high in cortical GABAergic neurons as shown ^50^. With VGAT-Cre mediated deletion of *Cnr1* (*VGAT*^Cre^:*Rosa26*^EYFP^:*Cnr1*^fl/fl^) these high *Cnr1* expressing cells were absent and there was no eYFP-*Cnr1* co-expressing neurons (Fig. 2A). Because CB1Rs are implicated in development of cortical circuits ^51, 52^, we used a heterozygous deletion strategy to reduce CB1Rs in GABAergic neurons in *Fmr1* KO mice (*VGAT^Cre^:Cnr1^fl/+^:Fmr1* KO) in order to decrease any confounds induced by developmental changes. Heterozygous *Cnr1* expression results in half the CB1R protein ^21^.

**Figure 2:**
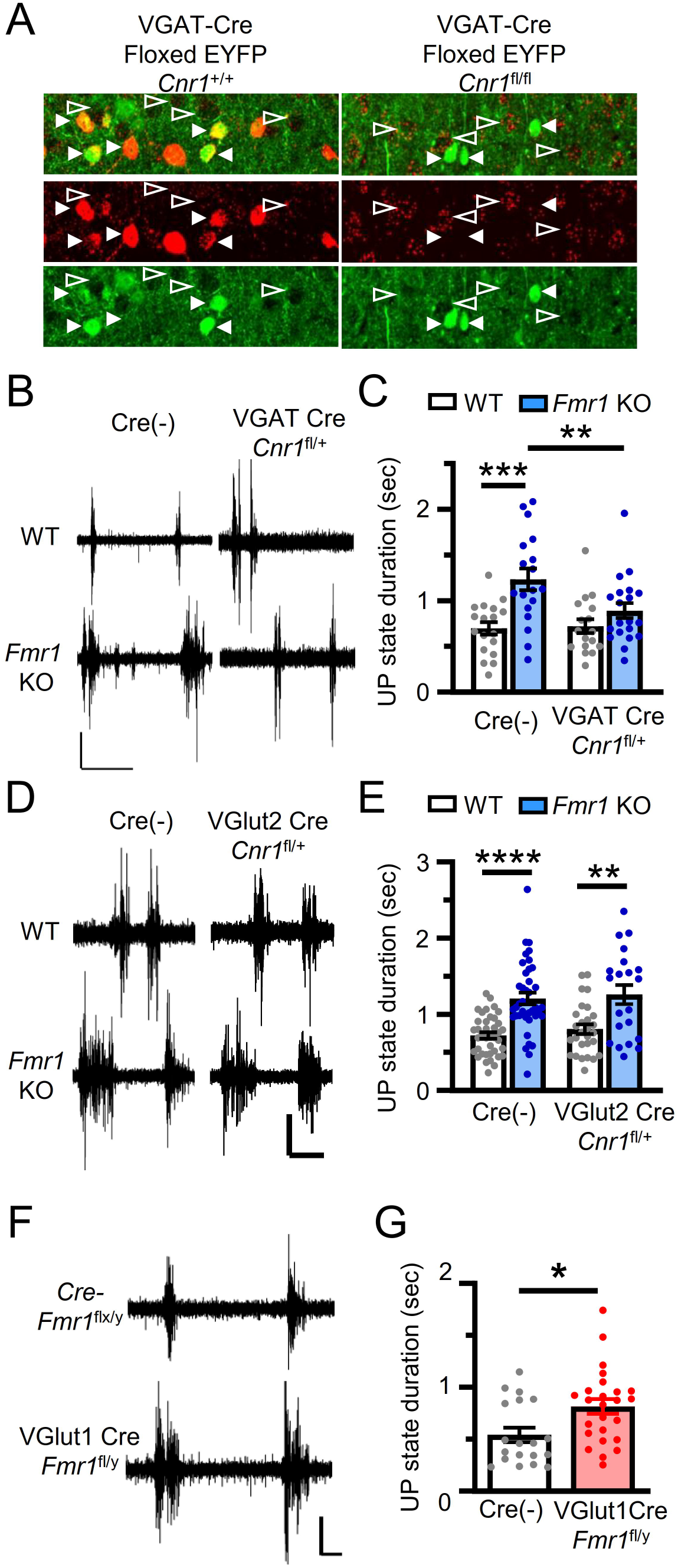
Genetic reduction of *Cnr1* in GABAergic neurons, but not glutamatergic neurons, rescues prolonged Up states in the *Fmr1* KO. A) *Cnr1* gene deletion in GABAergic neurons was accomplished by crossing the VGAT-Cre and floxed-*Cnr1* mouse lines. Images of homozygous deletion using IHC for the reporter EYFP (Green) and RNAscope for *Cnr1* message (Red) show our ability to effectively target gene dosage in GABAergic neurons. B,C) Heterozygous deletion of *Cnr1* in GABAergic neurons (VGATCre *Cnr1*^fl/+^) rescues Up state duration in *Fmr1* KO slices. B) Example traces of the four genotypic groups C) Group average (±SEM) of Up state duration (N=17-20 slices and 5-6 mice for each genotype). D,E) Heterozygous deletion of *Cnr1* in glutamatergic neurons (VGlut2Cre *Cnr1*^fl/+^) did not affect Up state duration in WT or *Fmr1* KO (N=21-39 slices and 5-9 mice for each genotype). F,G) Deletion of *Fmr1* in glutamatergic neurons is sufficient to cause long Up states. Floxed (fl) *Fmr1* mice were crossed to VGlut1-Cre mice to delete *Fmr1* in cortical glutamatergic neurons (VGlut1Cre *Fmr1*^fl/y^). F) Example traces from each genotype. G) Scatterplot and bar-graph showing mean duration is increased with glutamatergic neuron-specific *Fmr1* deletion (N=19,25 slices and 5,6 mouse pairs for each genotype). Scale bars for all panels = 50 µV and 1 s. *p<0.05; **p<0.01; ***p<0.005. 2-way ANOVA with posthoc comparisons.

If CB1R-mediated, presynaptic suppression at GABAergic synaptic transmission is enhanced in *Fmr1* KO mice, we would expect reduced *Cnr1* dosage in GABAergic neurons to increase inhibition and consequently, reduce and normalize Up state duration in *Fmr1* KO slices. In Cre(-) controls, with wildtype levels of *Cnr1* in GABAergic neurons, we reproduced the long Up state phenotype in *Fmr1* KO mice (Fig. 2B,C). Heterozygous deletion of *Cnr1* selectively in GABAergic neurons (*VGAT^Cre^:Cnr1^fl/+^: Fmr1* KO) reduced Up state duration as compared to Cre (-) *Fmr1* KO and durations were similar to Cre(-) WT. (Fig. 2B,C). These results are consistent with enhanced CB1R-mediated signaling in GABAergic causing longer Up states. Interestingly, reduction of *Cnr1* in GABAergic neurons on a *Fmr1* WT background (*VGAT^Cre^:Cnr1^fl/+^:* WT) had no effect on Up state duration suggesting that CB1R-mediated suppression of GABAergic transmission does not normally contribute to Up state duration in this context. Up state amplitude was reduced in *VGAT^Cre^:Cnr1^fl/+^* mice on both the WT and *Fmr1* KO background (2way ANOVA, Main effect of *Cnr1;* F (1, 69) = 9.150; p< 0.01; Fig. S2A,B), similar to what we observed with rimonabant (Fig. S1A).

CB1R-mediated, presynaptic suppression of glutamatergic synaptic transmission is decreased in the frontal cortex of *Fmr1* KO mice which may contribute to cortical circuit hyperexcitability ^17^. To examine the contribution of reduced CB1R-function in excitatory neurons to Up states in *Fmr1* KO neurons, we used the same strategy as above except we used VGlut2-Cre mice to delete *Cnr1* in cortical glutamatergic neurons ^38^.

Deletion of *Cnr1* in glutamatergic neurons was confirmed using RNAscope (Fig. S2C). To examine effects on Up states, we again used a heterozygous deletion strategy (*VGlut2^Cre^:Cnr1^fl/+^)* to reduce *Cnr1*. If reduced *Cnr1* function in excitatory neurons contributes to long Up states in the *Fmr1* KO, then we may expect Up states in *VGlut2^Cre^:Cnr1^fl/+^: Fmr1 ^-/y^* mice to be longer than in Cre(-) *Fmr1* KO slices. In contrast to this prediction, a reduction of *Cnr1* in glutamatergic neurons had no effect on Up state duration on either a WT or *Fmr1* KO background (Fig. 2D,E). *VGlut2^Cre^:Cnr1^fl/+^*mice also had normal Up state amplitude and frequency on both the WT or *Fmr1* KO backgrounds (Fig. S2D,E). These results suggest that reduced CB1R function in glutamatergic neurons does not contribute to long Up states in *Fmr1* KO mice and supports a role for enhanced CB1R function specifically in inhibitory neurons.

### Deletion of Fmr1 in glutamatergic neurons is sufficient to cause long Up states

Although our results implicate CB1R function in inhibitory neurons, we hypothesize that loss of FMRP in glutamatergic neurons leads to increased synthesis or release of endocannabinoids near GABAergic presynaptic terminals to suppress release. In support of this idea, our previous study found that simultaneous deletion of *Fmr1* in glutamatergic neurons and glia, but not deletion restricted to GABAergic neurons, recapitulated the long Up state duration phenotype ^16, 53^. Subsequent work implicated *Fmr1* deletion in astrocytes in circuit hyperexcitability, including long Up states ^54^. To re-address this issue, we created mice with conditional deletion of *Fmr1* in forebrain glutamatergic neurons (*VGlut1^Cre^:Fmr1 ^fl/y^*) using VGlut1-Cre ^37^ and floxed *Fmr1* (*Fmr1 ^fl/y^*) mice and measured Up state in comparison to Cre(-) male littermates. We found that *Fmr1* deletion in glutamatergic neurons was sufficient to induce longer Up states and mimic the *Fmr1* KO phenotype (Fig. 2F,G). Up state frequency and amplitude in *VGlut1^Cre^:Fmr1 ^fl/y^* slices were not different from Cre(-) littermates (Fig. S2F,G). Therefore, our results support the hypothesis that postsynaptic loss of *Fmr1* in glutamatergic neurons leads to enhanced CB1R-mediated presynaptic suppression of inhibitory synapses.

### Reduced Cnr1 dosage and CB1R antagonism rescue auditory-driven synchronous cortical activity in Fmr1 KO mice

Our results in slices support a role for enhanced CB1R function in GABAergic neurons underlying altered cortical network activity in the *Fmr1* KO. We next utilized EEG recordings to determine if enhanced CB1R function in GABAergic neurons contributed to altered resting and sensory-driven cortical activity *in vivo* in the *Fmr1* KO. Importantly, these EEG phenotypes are observed in individuals with FXS and results may inform mechanism and therapeutic targets for human EEG alterations ^5, 7, 10^. EEG measurements using a 30 point array of surface electrodes contacting the skull were performed as previously described ^8^. The array was divided up into 3 regions in each hemisphere (Frontal, Medial and Temporal) (Fig. 3A). We measured the ability of auditory stimuli to consistently drive synchronized cortical activity among all stimulus trials of sound presentation using intertrial phase coherence (ITPC) – a measure that focuses on the timing of frequency components independent of raw power.

**Figure 3.**
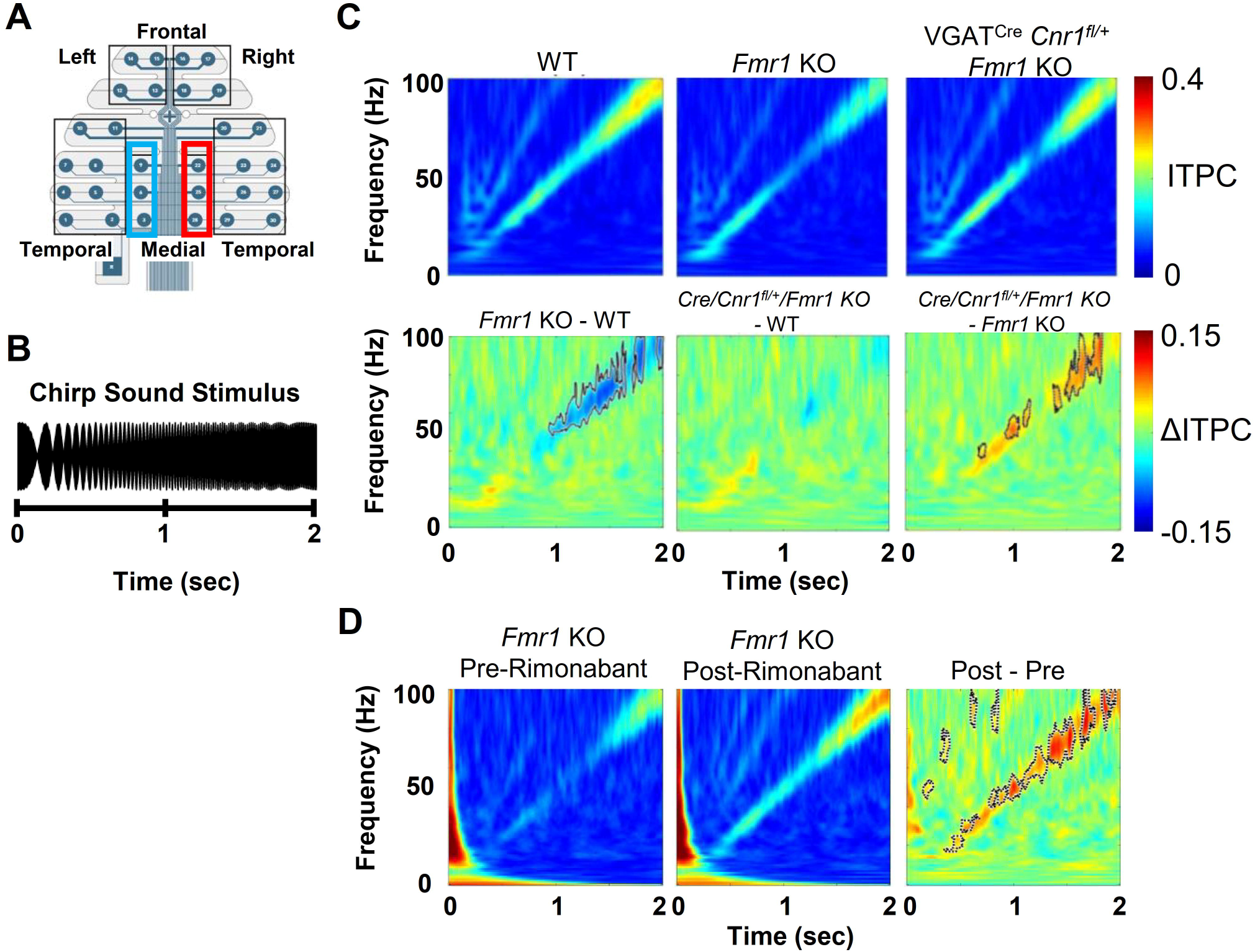
Genetic reduction of *Cnr1* in GABAergic neurons or rimonabant treatment rescue the decreased synchronization of chirp-driven cortical activity in the *Fmr1* KO. A) Diagram of EEG electrode array laid over scalp of mice. B) Chirp stimuli were 2 seconds of broadband noise sinusoidally modulated starting at 1 Hz and ramping up to 100 Hz. C) *Cnr1* genetic reduction experiment. *Top*: ITPC average plots of induced activity in the right medial region (red box in A) in the 3 genotypes examined (WT, *Fmr1* KO, and *Fmr1* KO with heterozygous deletion of *Cnr1* in GABAergic neurons; VGAT^Cre^/*Cnr1^fl/+^*/*Fmr1* KO). *Bottom*: Average difference plots (ΔITPC) based on plots above indicate decreased ITPC at higher frequencies in *Fmr1* KO mice (left) and the rescue of this phenotype with a genetic reduction of *Cnr1* in GABAergic neurons . N=17,17,17 mice. D) Rimonabant treatment experiment. *Left*: ITPC average plots of induced activity for pre- and post-rimonabant treatment obtained from the left medial region (blue box in A) in *Fmr1* KO mice (N=14). *Right*: Difference plot indicating increases in ITPC with rimonabant treatment. Solid lines border statistically different areas of the plot. Scales in C apply to D.

We first examined ITPC of activity induced by a chirp sound (Fig 5B). We performed the same *Cnr1* gene dosage reduction in GABAergic neurons and compared ITPC among littermates of these 3 genotypes: Cre (-) “WT”, Cre(-) *Fmr1* KO, and *Fmr1* KO with heterozygous deletion of *Cnr1* in GABAergic neurons (*VGAT^Cre^:Cnr1^fl/+^:Fmr1* KO). We focus first on the result averaged from the right medial region of the electrode array (Fig. 5A, red box). The average ITPC plots have a notable diagonal of higher ITPC that corresponds to modulation at the frequency currently occurring in the chirp (Fig. 3C, *top*). From the ITPC plots, average difference plots with plot areas of statistical difference marked by bold black line were calculated for each of the three possible genotypic comparisons (Fig. 3C, *bottom*). As reported previously for both FXS individuals and *Fmr1* KO mice, the “*Fmr1* KO – WT” difference plot revealed a decrease in ITPC in the higher frequency components corresponding to the frequency modulation occurring later in the chirp (Fig. 3C, *bottom left*) ^6, 7^ ^8^. This phenotype was observed in all 6 electrode array regions (Fig. S3). This decrease indicated an impaired ability for higher frequency sound modulation to consistently drive cortical synchrony from one chirp trial to the next.

The two other difference plots (Fig. 3C, *bottom*) indicate that the decrease in *Cnr1* dosage rescued the phenotype. First, the “*VGAT^Cre^:Cnr1^fl/+^:Fmr1* KO– WT” difference plot (Fig. 3C, *bottom middle*) did not have any regions of difference indicating that reduced *Cnr1* dosage rescued the decreased ITPC in the *Fmr1* KO mouse. Second, consistent with the changes just described, the “*VGAT^Cre^:Cnr1^fl/+^:Fmr1* KO– *Fmr1* KO” difference plot (Fig. 3C, *bottom right*) indicates increased ITPC in *Fmr1* KO mice with decreased *Cnr1* dosage. Therefore, decreased *Cnr1* dosage in GABAergic neurons increased the ability of sounds to consistently drive cortical activity in *Fmr1* KO mice. Interestingly, this rescue occurred in both medial regions of the electrode array but did not occur in frontal or temporal regions (Fig. S3).

Acute pharmacological blockade of CB1R rescues Up state duration in *Fmr1* KO slices (Fig. 1). We next determined if rimonabant treatment could correct the ITPC in the *Fmr1* KO In addition, to determine the potential of chronic CB1R antagonism as a therapeutic strategy, we tested a 7-day treatment with the CBR1 antagonist. We recorded pre- and post-treatment from *Fmr1* KO mice. Rimonabant increased ITPC in the *Fmr1* KO, which is consistent with a rescue to wild-type levels (Fig. 3D). Unlike gene dosage, this rescue occurred in all regions of the electrode array (Fig. S4).

We obtained similar results examining ITPC of an auditory steady state response (ASSR). The ASSR was induced with either a 40 Hz or 80 Hz click train. The clearest data were collected with the 80 Hz ASSR, but similar results were also observed with 40 Hz ASSR (Figs. 4,5, S5-S8). There was a robust decrease in ITPC in the 80 Hz band in Cre (-) *Fmr1* KO compared to Cre (-) WT as seen in the difference plot (Fig. 4C, *bottom left*). The ITPC difference was non-existent when comparing *VGAT^Cre^:Cnr1^fl/+^:Fmr1* KO to WT (Fig. 4C, *bottom middle*) and ITPC in the *VGAT^Cre^:Cnr1^fl/+^:Fmr1* KO mice was greater than that observed in *Fmr1* KO littermates (Fig. 4C, *bottom right*). The 80 Hz ASSR decrease in the *Fmr1* KO mice was seen and rescued by heterozygous deletion of *Cnr1* in GABAergic neurons in all brain regions measured with the electrode array (Fig. S5). Similar results were observed with ITPC of a 40 Hz ASSR (Figs. 5C; S7). Like that observed with genetic reduction of CB1Rs in GABAergic neurons, a 7-day rimonabant treatment of *Fmr1* KO mice increased ITPC for 40 and 80 Hz ASSR (Figs. 4D, 5D, S6, S8) suggesting that rimonabant treatment could also rescue normal auditory-driven synchrony.

**Figure 4.**
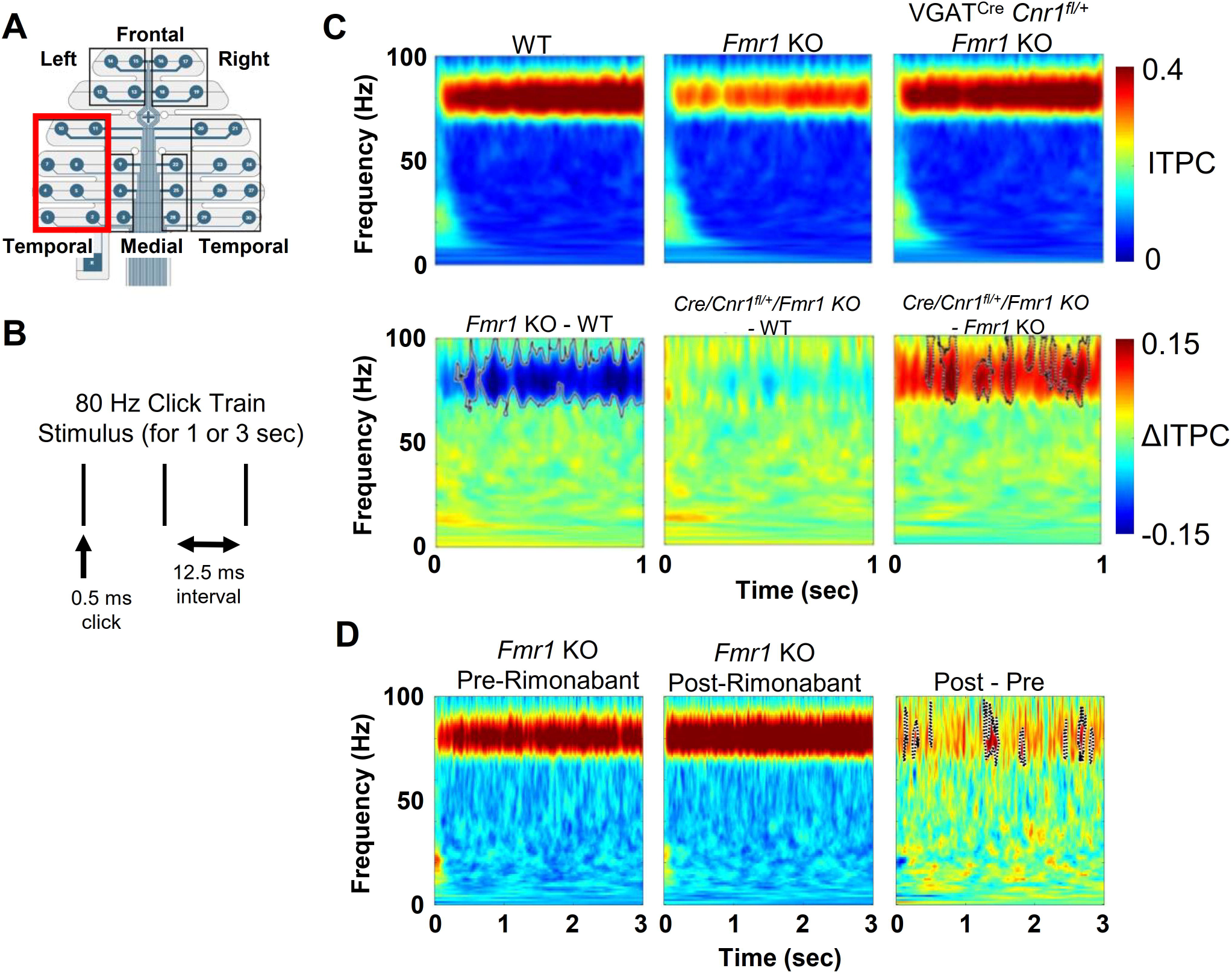
Both genetic reduction of *Cnr1* in GABAergic neurons and rimonabant rescue decreased synchronization of pulse-driven cortical activity at 80 Hz in the *Fmr1* KO. A) Diagram of EEG electrode array. Data shown are from the left temporal region electrodes (red box). B) An 80 Hz pulse train was used to induce an ASSR (auditory steady-state response). C) *Top*: ITPC average plots of induced activity in the 3 genotypes examined. *Bottom*: Average difference plots indicate decreased ITPC during the ASSR in *Fmr1* KO mice (left) and the rescue of this phenotype in *Fmr1* KO mice with genetic reduction of *Cnr1* in GABAergic neurons (VGAT^Cre^/*Cnr1^fl/+^*/*Fmr1* KO). N=17,17,17 mice. D) *Left*: ITPC average plots of induced activity for pre- and post-rimonabant treatment in *Fmr1* KO mice (N=14). *Right*: Difference plot indicating an increase in ITPC with rimonabant treatment. Solid lines border statistically different areas of the plot. Scales in C apply to D.

**Figure 5.**
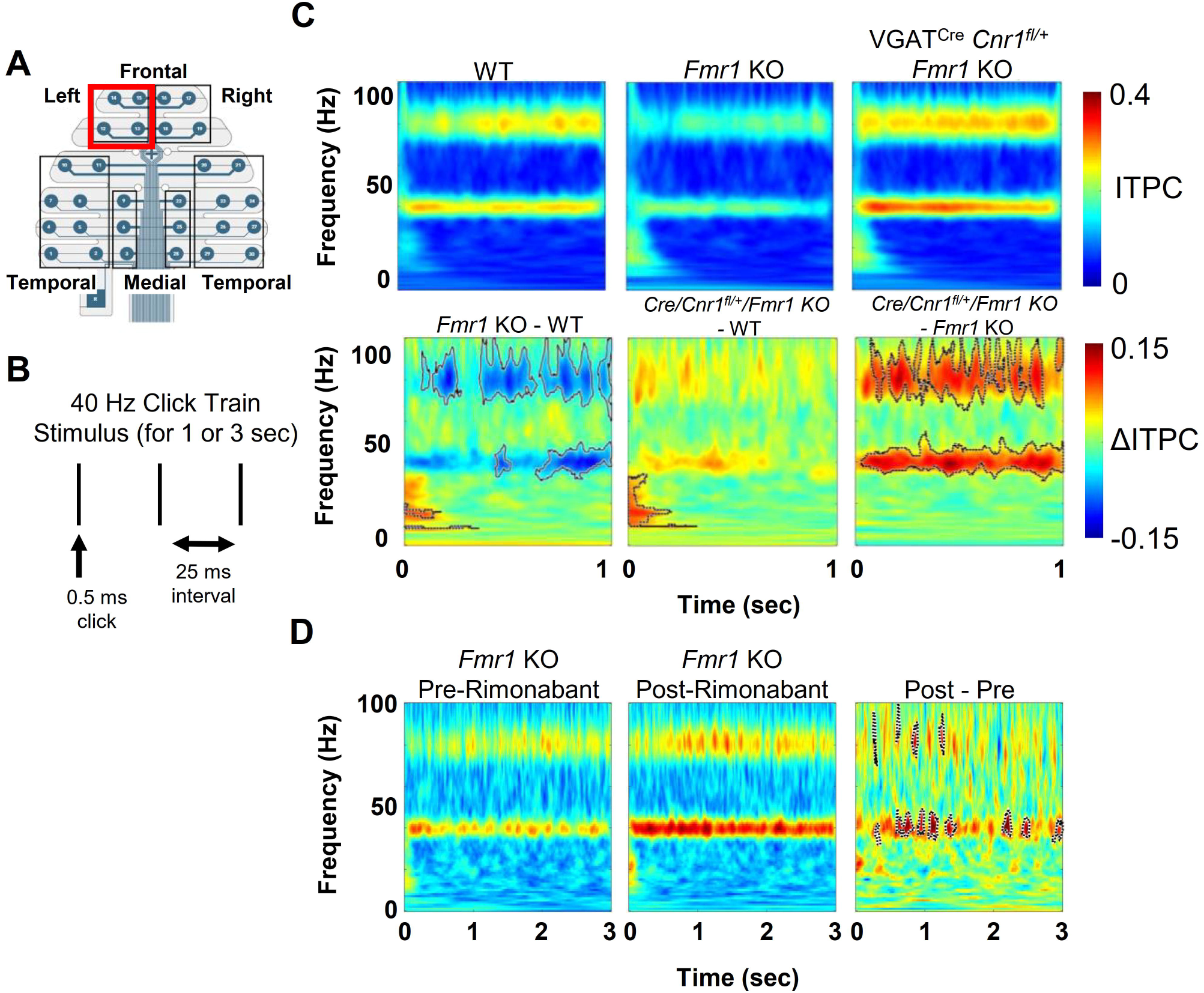
Both genetic reduction of *Cnr1* in GABAergic neurons and rimonabant rescue decreased synchronization of pulse-driven cortical activity at 40 Hz in the *Fmr1* KO. A) Diagram of EEG electrode array. Data shown are from the left frontal region electrodes (red box). B) A 40 Hz pulse train was used to induce an ASSR. C) Genetic reduction experiment. *Top*: ITPC average plots of induced activity in the 3 genotypes examined. *Bottom*: Average difference plots indicate decreased ITPC during the ASSR in *Fmr1* KO mice (left) and the rescue of this phenotype in *Fmr1* KO with genetic reduction of *Cnr1* in GABAergic neurons (VGAT^Cre^/*Cnr1^fl/+^*/*Fmr1* KO). N=17,17,17 mice. D) Rimonabant treatment experiment. *Left*: ITPC average plots of induced activity for pre- and post-rimonabant treatment obtained from the left medial region in *Fmr1* KO mice (N=14). *Right*: Difference plot indicating increases in ITPC with rimonabant treatment. Solid lines border statistically different areas of the plot. Scales in C apply to D.

**Figure 6.**
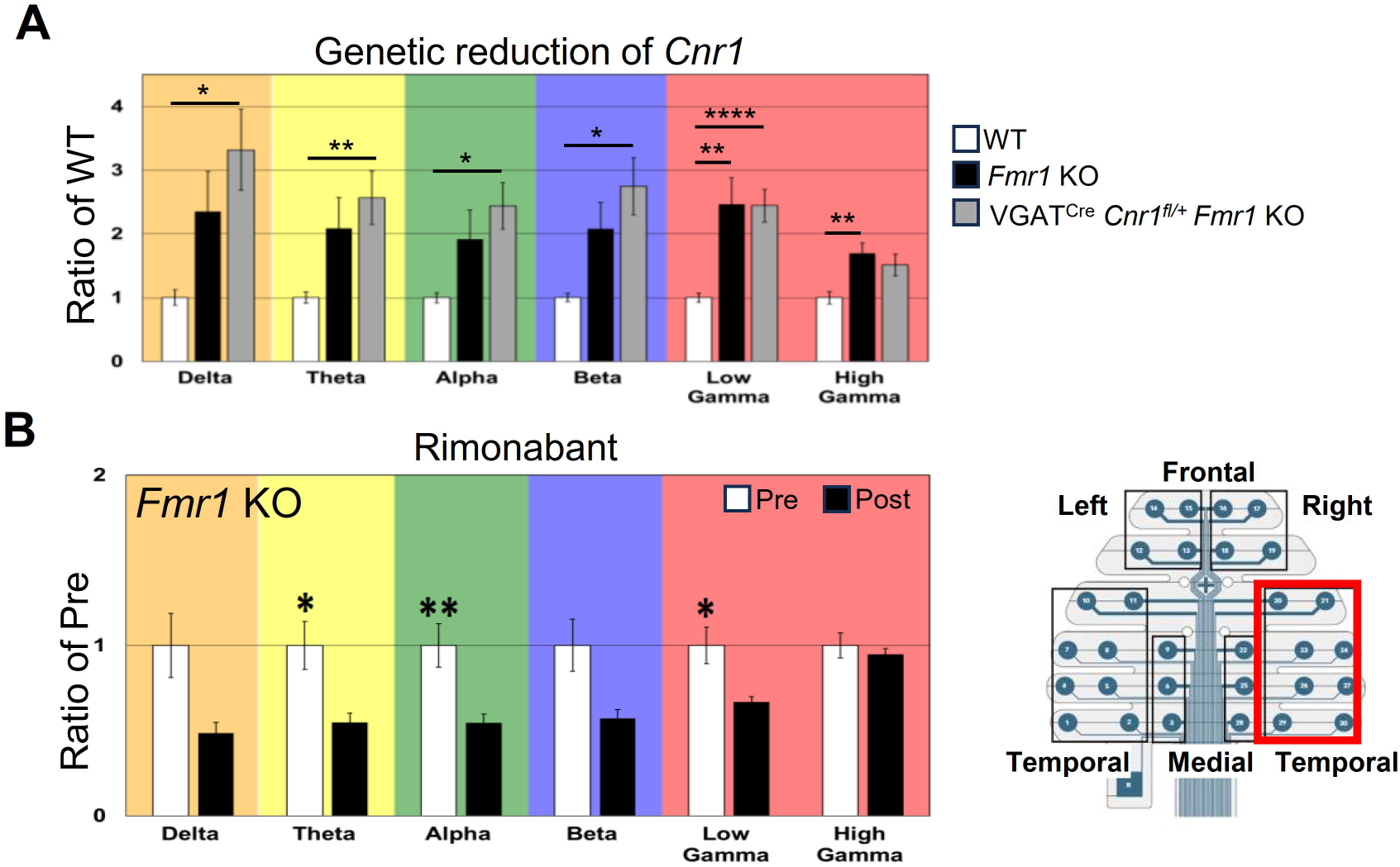
Distinct effects of genetic and pharmacological reduction of CB1R activity on resting state EEG. Spectral density power (normalized to wild-type (WT) mice (A) or to pre-treatment *Fmr1* KO mice (B)) is plotted for 6 standard frequency bands and obtained from the right temporal electrode group (as indicated by red box in lower right). A) An increase in power is observed in the low and high gamma frequency bands in the *Fmr1* KO mice (*black*) and as compared to WT littermates. *Fmr1* KO mice with genetic reduction of *Cnr1* in GABAergic neurons (VGAT Cre/*Cnr1*^fl/+^*/Fmr1* KO) (gray) have increased power across all frequency bands, as compared to WT, and is not different from *Fmr1* KO, with the exception of high gamma. N=17, 17, 17 mice. B) 7 day rimonabant treatment of *Fmr1* KO mice decreases power in all frequency bands except High Gamma. *, p<0.05 **, p<0.01 ****, p<0.001

In summary, these data indicate that enhanced CB1R signaling at GABAergic synapses is involved in systems level auditory response changes in the *Fmr1* KO mouse. Also, a pharmacological block of CBR1 later in life is sufficient to rescue these deficits.

### CBR1 antagonism may rescue resting activity in vivo, but reduced Cnr1 dosage in GABAergic neurons does not

Under resting conditions when no experimental and salient stimuli are being presented, power in the standard EEG frequency bands – and gamma in particular - is higher in both FXS individuals and in the *Fmr1* KO mouse ^5, 6, 55^. We determined if reduced *Cnr1* dosage in GABAergic neurons could rescue this resting power phenotype. For most electrode array regions, increased power was observed in the *Fmr1* KO across all frequency bands when compared to WT controls, but most consistently in the lower gamma band (Fig. 6A, S9). But there was no clear example of rescue in any of the standard frequency bands. Almost all the average power values measured in *VGAT^Cre^:Cnr1^fl/+^:Fmr1* KO mice were different from those in WT mice, but not *Fmr1* KO mice, indicating that heterozygous deletion of *Cnr1* in GABAergic neurons did not rescue the resting state power phenotypes in the *Fmr1* KO.

We next examined the effect of 7-day rimonabant treatment on resting power observed in the *Fmr1* KO (Figs. 6B, S10). Unlike the reduced *Cnr1* dosage, rimonabant clearly decreased the power of all frequency bands except high gamma, and this change was consistent with an ability to rescue the power phenotypes towards WT levels. This pattern was observed at all electrode regions. While the data suggest that altered, ongoing CBR1 signaling is involved in resting power changes in the *Fmr1* KO mice, this signaling is more effectively remedied pharmacologically in adults, and not by cell type specific genetic reduction throughout life.

## Discussion

Our results provide evidence for enhanced, cell-type specific eCB signaling underlying cortical network function phenotypes in *Fmr1* KO mice. Specifically, our findings support the hypothesis that enhanced CB1R-mediated suppression of GABAergic synaptic transmission mediates prolonged cortical Up states and deficits in sensory-driven cortical network synchrony in the *Fmr1* KO mouse. This conclusion is supported by the “rescue” (or normalization) of circuit activity and EEG phenotypes using pharmacological blockade of CB1Rs or 2-AG synthesis and *Cnr1* gene dosage reduction in GABAergic neurons. We also observe that GABAergic synaptic currents in *Fmr1* KO L2/3 cortical neurons are more sensitive to antagonism of CBR1 receptors consistent with enhanced CB1R-suppression of GABAergic transmission. While selective *Cnr1* deletion in glutamatergic neurons has no effect on Up states, *Fmr1* deletion in the same neurons is sufficient to mimic the long Up state phenotype observed in constitutive *Fmr1* KO. Taken together with previous work, our results support a model where loss of FMRP in cortical glutamatergic neurons leads to altered or enhanced mGluR5-dependent synthesis of 2-AG and enhanced suppression of GABA release from CB1R-positive inhibitory presynaptic terminals. Importantly, our results implicate a specific synaptic mechanism in controlling the duration of cortical Up states as well as the ability of cortical circuits to appropriately synchronize with time-modulated stimuli in the gamma frequency range, as measured with ITPC. Because sensory-driven cortical synchrony and ITPC are deficient in individuals with FXS, our results suggest that enhanced eCB function in cortical circuits may contribute to abnormal temporal processing of sensory stimuli in FXS.

### Altered modulation of CB1R-regulated inhibitory circuits as a mediator of prolonged circuit activity in FXS

Up states are mediated by local cortical excitatory and inhibitory circuits and thus reflect the balance of excitation and inhibition ^56^. If or how inhibitory neurons control the termination of Up states or their duration has been well studied in wildtype animals^57^. Interestingly, pharmacological blockade of GABAa receptors reduces Up state duration, whereas blockade of GABAB receptors (GABAbRs) increases Up state duration ^58, 59^. This suggests that CB1R-mediated suppression of GABABR activation may contribute to Up state duration in *Fmr1* KO ^60^. Consistent with this idea, the GABAB agonist baclofen increases cortical synchronization (chirp ITPC) in the *Fmr1* KO ^61^. Opto- or chemogenetic inhibition of specific inhibitory neuron types, such as parvalbumin and somatostatin-positive interneurons, enhance pyramidal neuron firing during Up states and prolong Up state duration both *in vivo* and in slices ^62, 63^. Because CB1Rs are highly expressed on presynaptic terminals of Cholecystokinin-positive inhibitory interneurons ^64^, it is likely that dysregulation of input from CCK-interneurons underlie prolong Up state duration in *Fmr1* KO. The role of CCK-interneurons in Up state regulation in wildtype cortex is unknown, perhaps due to the difficulty in genetically targeting this cell type in cortex ^65^. We did not observe effects of pharmacological or genetic reduction of CB1R activity on Up state duration in mice that were WT for *Fmr1,* indicating that CB1Rs and perhaps CCK+ mediated inhibition do not regulate Up state duration in normal developing cortex as reported previously^41^. Therefore, our results indicate that CB1Rs are abnormally active during Up states in the cortex of young *Fmr1* KO mice under our experimental conditions.

Although CB1R-dependent suppression of excitatory synaptic transmission is reportedly reduced in *Fmr1* KO cortex ^17^, genetic reduction of CB1Rs in cortical glutamatergic neurons did not mimic or exacerbate the long Up states in the *Fmr1* KO. This result suggests that the primary effects of CB1Rs on Up states in the *Fmr1* KO are via regulation of inhibitory synaptic transmission. CB1Rs are highly concentrated on presynaptic terminals of CCK+ inhibitory neurons^50, 64^. Because we observe strong potentiation of inhibitory synaptic currents with rimonabant treatment of *Fmr1* KO slices in the absence of action potentials (in TTX), there may be enhanced tonic CB1R-dependent suppression of inhibitory synaptic transmission in the *Fmr1* KO ^60, 66, 67^. Because the DAGLα inhibitor, DO34, rescues Up state duration in *Fmr1* KO slices, there may be enhanced 2-AG synthesis in *Fmr1* KO excitatory neurons that tonically suppress GABAergic transmission. While we observed a enhanced suppression of IPSCs by eCB signaling in *Fmr1* KO slices, this was not observed in a previous study in hippocampus^21^ and could be due to different experimental conditions, or brain regions.

### Molecular basis for enhanced CB1R-dependent suppression of inhibition in Fmr1 KO

mGluR5 signaling contributes to prolonged Up states^16^ and stimulates 2-AG synthesis through activation of phospholipase Cβ and DAGLα ^42^. Because mGluR5-induced and CB1R-dependent suppression of inhibitory synaptic transmission is enhanced in the striatum and hippocampus of *Fmr1* KO mice^18–20^, we hypothesize that this is the case in the neocortex. In *Fmr1* KO mice, CB1R levels are normal and CBR1 agonist-induced suppression of inhibition is normal^18, 20^. These results argue against upregulation of CB1R function in the *Fmr1* KO but enhanced, and perhaps tonic, mGluR5- dependent 2AG synthesis selectively at inhibitory synapses. In support of this idea, basal 2-AG levels are elevated or saturated in *Fmr1* KO striatum and cortex, and insensitive to the stimulatory effects of mGluR5 agonism^17, 19^. A molecular basis for enhanced mGluR5 and CB1R-dependent suppression of inhibitory synaptic transmission in the *Fmr1* KO may be related to mislocalization of DAGLα and mGluR5 away from the postsynaptic density^17, 68^, which may enhance their concentration and function near CCK+ GABAergic terminals with high levels of CB1Rs ^69^.

### Enhanced CB1R function in inhibitory neurons mediates deficits in sensory-driven cortical synchronization and coherence in awake Fmr1 KO mice

Here we demonstrate that the pharmacological and genetic manipulations of CB1Rs that correct Up state duration also correct some EEG phenotypes that are conserved in FXS individuals. Cortical synchronization (or ITPC) during the chirp, 40 and 80 Hz ASSR in the *Fmr1* KO was strongly and consistently increased by both rimonabant and heterozygous deletion of *Cnr1* in inhibitory neurons. This result suggests that the local cortical circuit mechanisms that mediate prolonged Up states in the *Fmr1* KO also contribute to an inability of cortical circuits to consistently synchronize with rapidly modulated sensory stimuli. Cannabinoids are known to regulate neural synchrony and gamma oscillations; studied in the context of the effects of cannabis use on cognition or in schizophrenia^70, 71^. Specifically, intravenous application of Δ-9-tetrahydrocanabinol (Δ9-THC) in humans, the primary psychoactive constituent of marijuana and a CB1R agonist, reduced evoked power during the 40 Hz ASSR as well as ITPC ^72^. This result is consistent with the idea that overactive CB1R signaling *Fmr1* KO mice leads to reduced ASSR and reduced cortical synchronization and may function similarly in FXS individuals.

Although ASSR ITPC deficits were rescued across all brain regions by *Cnr1* deletion in GABAergic neurons, the chirp ITPC deficits were rescued in only medial cortical regions. There are reported cortical region differences in expression of CCK+ inhibitory interneurons which may contribute differently to the chirp ITPC ^73^.

Similarly, frontal cortical areas have higher levels of CB1Rs as compared to primary sensory cortex and genetic reduction may have less of a functional impact in frontal areas^74^. Alternatively, because there is a known decreased function of CB1Rs at excitatory synapses in the frontal cortex, this may make a larger contribution to the frontal cortex ITPC deficits in the *Fmr1* KO^17^. We also observed distinct effects of pharmacological vs genetic reduction of CB1R activity on resting state EEG. While rimonabant reduced resting state gamma power in *Fmr1* KO mice, genetic reduction of *Cnr1* in inhibitory neurons did not. This may be due to differences in the effects of CB1R antagonism in adult vs throughout development ^72, 75^. Alternatively, antagonism of CB1Rs on excitatory neurons by rimonabant may mediate effects on resting state EEG.

### ASD risk genes regulate E/I balance through CB1Rs

In addition to *Fmr1* , loss of function of other ASD risk genes such as Neuroligin3 or Neurexin 1B affect CB1R-regulation of synaptic transmission^49, 76, 77^. With *Nlgn3* mutations, tonic CB1R-mediated suppression of IPSCs is deficient whereas loss of function of *Nrxn1B* enhances tonic CB1R suppression of excitatory synaptic transmission. These results reveal how ASD risk genes can affect the E/I balance but do so through different synaptic mechanisms. Based on the known role of eCBs in regulation of neural oscillations^70, 71^ and our observations that we can ameliorate deficits in sensory-driven network oscillations in *Fmr1* KO mice by manipulation of CB1Rs, dysfunction of CB1Rs may contribute to altered neural oscillations and deficits in sensory processing and cognition in these mouse or individuals with loss of function in these other ASD risk genes. Furthermore, our results suggest that targeting selective CB1R antagonism may be an effective therapeutic strategy to restore cortical synchronization and aid sensory processing in individuals with FXS.

## Acknowledgements

We would like to acknowledge Jacob E. Bowles and Christopher Williams for technical assistance with genotyping and Dr. Laurent Gautron for assistance with RNAscope. This work was supported by NIH grants U54HD082008 and U54HD104461 (KMH, JRG, DKB) and P30DK127984 (UTSW Metabolic Core).

