## Supplemental methods and figures for "Enhanced CB1 receptor function in GABAergic neurons mediates hyperexcitability and impaired sensory-driven synchrony of cortical circuits in Fragile X Syndrome model mice"

**Supplementary Information:**

**Supplementary Methods:**

*Acute brain slice preparation:* Slice preparation was based on our earlier study [1]. Male mice (18 – 24 days of age) were deeply anesthetized with ketamine/xylazine cocktail (4 mg/ml xylazine in 30 mg/ml ketamine)) and decapitated. The brain was transferred into an ice-cold dissection buffer containing (in mM): 87 NaCl, 3 KCl, 1.25 $NaH_2PO_4$, 26 $NaHCO_3$, 7 $MgCl_2$, 0.5 $CaCl_2$, 20 D-glucose, 75 sucrose, and 1.3 ascorbic acid. The buffer was constantly aerated with 95% $O_2$-5% $CO_2$ gas cocktail. The brain was glued to an angled block [2]. Thalamocortical slices, 400 μm, were cut using a vibratome (Leica VT 1200S). Slices were immediately transferred to an interface recording chamber (Harvard Instruments) and allowed to recover for 1 h in nominal artificial CSF (ACSF) at 32°C containing (in mM): 126 NaCl, 3 KCl, 1.25 $NaH_2PO_4$, 26 $NaHCO_3$, 2 $MgCl_2$, 2 $CaCl_2$, and 25 D-glucose. After the recovery slices were perfused with a modified ACSF that better mimics physiological ionic concentrations *in vivo* [3] containing (in mM): 126 NaCl, 5 KCl, 1.25 $NaH_2PO_4$, 26 $NaHCO_3$, 1 $MgCl_2$, 1 $CaCl_2$, and 25 D-glucose (based on but modified from [1, 3]). Slices remained in this modified ACSF for 45 min followed by the UP state recordings.

*UP States recording and analysis:* Spontaneously generated UP states in brain slices were extracellularly recorded using 0.5 MΩ tungsten microelectrodes (FHC) placed in layer 4 of primary somatosensory cortex. Only 1 recording was obtained from each slice and data was collected from 4-8 slices per mouse. Slices were excluded or not recorded from if <2 UP states occurred per minute. This extracellular monitoring of UP states is a reliable indicator of the synchronous, depolarized state of neuron populations from which the term "UP state" was originally defined (Sanchez-Vives and McCormick, 2000; Rigas and Castro-Alamancos, 2007). A total of 5 min of spontaneous activity was collected from each slice. Recordings were amplified 10,000-fold, sampled at 2.5 kHz, and filtered on-line between 500 Hz and 3 kHz. All measurements were analyzed off-line using custom Labview software. For visualization and analysis of UP states, traces were offset to zero, rectified, and low-pass filtered with a 0.2 Hz cutoff frequency. Using these processed traces, the threshold for detection was set at 15× the RMS (root mean square) noise. An event was defined as an UP state if its amplitude remained above the threshold for at least 200 ms. The end of the UP state was determined when the amplitude decreased below threshold for >600 ms. Two events occurring within 600 ms of one another were considered a single UP state. UP state amplitude was defined based on the filtered/rectified traces and was unitless since it was normalized to the detection threshold. This amplitude may be considered a coarse indicator of the underlying firing rates of neuronal populations. Direct measures of firing rates were not possible because individual spikes could not be isolated except during the quiet periods (the DOWN states).

*Drug application to slices:* Rimonabant (SR141716A, 5 μM, Tocris, Cat 0923. 0.5% DMSO) was used to block CB1Rs. DO34 (10 μM, Glixx Laboratories Inc., 0.5% DMSO) was used to inhibit the synthetic enzyme for 2-arachidonoylglycerol (2-AG), diacylglycerol lipase-α (DAGL- α). Experiments employing Rimonabant and DO34 were performed in a paired comparison where half the slices from one mouse were used for vehicle and the other half, for drug. In this division, adjacent slices from the same mouse were placed into the two treatment groups. Compounds were applied immediately after slicing for 2 hours and recordings performed with the compounds present.

*MEA implantation:* Surgical and recording procedures were similar to those recently described [4]. Mice were anesthetized with isoflurane inhalation (0.2-0.5%) and given ketamine (80 mg/kg, i.p.) (Zoetis,10004027) and xylazine (10 mg/kg, i.p.) (Covetrus, 061035). Mice were aseptically prepared for surgery and secured in a stereotaxic apparatus. Artificial tear ointment was applied to the eyes to prevent drying. Toe pinch reflex was used to measure anesthetic depth throughout the surgery, and supplemental doses of ketamine/xylazine were administered as needed. Once the mouse was anesthetized, a midline sagittal incision was made along the scalp to expose the skull. A cotton-tip applicator was used to remove the periosteum from the skull and to clean skull with saline. A surgical marker was used to mark bregma and positions of three screws. A dental drill was used to drill 1mm diameter holes in the skull overlying the left frontal cortex, left cerebellum and right cerebellum. Screws (Protech International, 00-96 X 1/16) were advanced into drilled holes until secure; special care was taken not to advance the screws beyond the point of contact with the dura. Bregma was marked with a surgical marker and the probe grounding wire was placed in the nuchal musculature. Saline was added to the top of the skull to aid in probe adherence. The saline was allowed to dry, and Teflon was placed on top of the probe. Dental cement (Kuraray, 3382KA) was applied around the screws, on the base of the cotton-tip applicator post, and the Teflon covering the probe. Waterproof medical tape was used to secure the cotton-tip applicator to the probe connector. Triple antibiotic was applied along the edges of the dental cement followed by a subcutaneous injection of 0.1mg/kg buprenorphine (Reckitt & Colman, 5053624). Mice were placed on a heating pad to aid in recovery from anesthesia. Additional doses of buprenorphine were administered every 6-8 hours for continuous analgesia during the first 48 hours after surgery. EEG recordings were conducted 2-3 days after MEA implantation.

*EEG recording:* All EEG recordings were conducted in a sound-attenuated chamber lined with anechoic foam (Gretch-Ken Industries, Oregon). EEG recordings were obtained from awake and freely moving mice using the SmartBox (NeuroNexus) acquisition system. The acquisition hardware was set to lower (0.5 Hz) and upper (500 Hz) filters and data were sampled at a rate of 1250 Hz. Mice were connected to a headstage, under brief isoflurane anesthesia, and tethered by a SmartLink cable to a freely rotating commutator positioned directly above a plastic arena. The plastic arena was surrounded by a Faraday cage securely fixed to a vibration isolation table. Mice were allowed to habituate to the arena for 20 minutes before EEG recordings were obtained.

*Auditory stimulus presentation for EEG recordings*: Acoustic stimuli were generated using RPVDSEX software and RZ6 hardware (Tucker Davis Technologies, FL) and presented through a free-field speaker (MF1 Multi-Field Magnetic Speaker; Tucker-Davis Technologies, FL) located 12 inches directly above the arena. Sound pressure level (SPL) was modified using programmable attenuators in the RZ6 system. The speaker output was ~70dB SPL at the floor of the recording chamber with fluctuation of ± 3 dB for frequencies between 5 and 35 kHz as measured with a ¼ inch Bruel & Kjaer microphone. Sound delivery was synchronized with EEG recordings using a TTL pulse to mark the onset of each sound in a train.

After mice were habituated, EEG was recorded for 5 minutes in the absence of any specific auditory stimulation ('resting EEG'). Subsequently, acoustic stimulation was presented. To quantify the ability of neural generators to produce synchronized oscillations to time varying stimuli, we employed two types of auditory stimuli. The first type of stimulation is called the auditory chirp-modulated sound (henceforth, "chirp"). The chirp is a broadband noise stimulus whose amplitude is modulated using a sinusoid with increasing or decreasing frequency in the 1-100 Hz range [5-7]. The chirp facilitates a rapid measurement of evoked phase locking to auditory stimuli of varying frequencies and can be used to compare temporal processing in clinical and pre-clinical settings[7]. Inter-trial phase coherence (ITPC), also known as phase locking factor [8], can be used to determine the ability of neural generators to synchronize oscillations to the frequencies present across trials. Both humans with FXS and *Fmr1* KO mice show ITPC deficits in the gamma band frequencies (~40 Hz) [9-11]. In this study, each chirp stimulus was 2 seconds in duration, and the depth of modulation was 100%. Chirp trains were presented 200 times each with the interval between each train randomly generated to be between 1-1.5 s.

Second, we used a click train to assess the auditory steady-state response (ASSR). The ASSR has been used as a diagnostic biomarker for disorders such as schizophrenia [12, 13]. The ASSR drives steady brain oscillations at specific frequencies of interest. In this study, 40 and 80 Hz gamma frequencies were used to obtain ASSR, with the 40 Hz and 80 Hz generators likely located in cortex and brainstem, respectively [14, 15]. The ASSR stimulus trains consisted of 0.5 ms clicks repeated at a rate of either 40 or 80 Hz over a 1 or 3 s period. Each train was presented 50 times with an inter-train interval of 2 s.

For *Cnr1* reduced gene dosage EEG recordings, the chirp was preceded by a 1 second sound intensity ramp. The 40 and 80 Hz auditory steady state response (ASSR) was measured from 50 trials constituting a train of 0.5 ms clicks maintained for 1 second. The inter-trial interval was 2 seconds. Data were collected from three genotypic groups of 17 mice each: WT, *Fmr1* KO, and *Fmr1* KO/*Cnr1* heterozygous. For rimonabant EEG recordings, recordings were performed immediately before treatment and 2 h after the 7th day treatment of rimonabant (1mg/kg IP, once daily). This dose was the minimum effective amount reported in previous studies [16-18]. For the chirp auditory stimulus in these experiments, a 1 second sound intensity ramp preceding the chirp stimulus was not included. The ASSR was the same as for reduced gene dosage experiments except the click train was maintained for 3 seconds. Data were collected from 14 *Fmr1* KO mice.

*Statistical analyses for EEG:* All EEG files extracted from SmartBox software were saved in a format compatible with Analyzer 2.2 (Brain Vision Inc). Resting EEG recordings were first down sampled to 625 Hz and notch filtered at 60 Hz to remove any residual line frequency power. A semi-automatic procedure implemented in Analyzer 2.2 was used for artifact rejection after visual inspection of all EEG files. Less than 20% of data were rejected due to artifacts from any single animal recording.

Resting state data were divided into 1 s segments and each segment was subjected to Fast Fourier Transforms (FFT) analysis using a 10% Hanning window at 0.5 Hz bin resolution. The average power ($\mu V/Hz^2$) was calculated for each mouse from 1-100 Hz. Power was binned according to spectral frequency bands: Delta (1-4 Hz), Theta (4-8 Hz), Alpha (8-13 Hz), Beta (13-30 Hz), Low Gamma (30-55 Hz), and High Gamma (65-100 Hz).

Resting state EEG data were blinded and analyzed using two-way ANOVA with Genotype (WT, *Fmr1* KO) and Frequency (delta to gamma) for the cortical regions (left frontal, right frontal, left medial, right medial, left temporal and right temporal) as factors. Data were expressed as ratio of WT values to gauge relative differences in various factors using the same scale. Data were analyzed for each factor and corrected for using Bonferroni adjusted p-values. $p$ values < 0.05 were considered significant for ANOVA. In all cases where genotype means are reported, SEM was used. Statistical analyses were performed using the program GraphPad Prism 10.3.1.

Chirp and ASSR traces were processed with Morlet wavelets linearly spaced from 1-100 Hz using voltage (µV). Wavelet coefficients were exported as complex values for use with inter-trial phase coherence (ITPC) analysis. Wavelets were run with a Morlet parameter of 10. To measure phase synchronization at each frequency across trials, ITPC was calculated as follows:

$$ITPC(f,t) = \frac{1}{n}\sum_{k=1}^{n}\frac{F_k(f,t)}{|F_k(f,t)|}$$

where $f$ is the frequency, $t$ is the time point, and $k$ is trial number. Thus, $F_k(f,t)$ refers to the complex wavelet coefficient at a given frequency and time for the $k$th trial.

Statistical group comparisons of ITPC in chirp and ASSR (40 and 80Hz) traces were quantified using a Monte Carlo permutation approach. Analysis was conducted by binning time into 256 parts and frequency into 100 parts, resulting in a 100 x 256 matrix. Non-parametric analysis was used to determine contiguous regions in the matrix that were significantly different from a distribution of 2000 randomized Monte Carlo permutations based on previously published methods [19]. Cluster sizes of the real genotype (both positive and negative direction, resulting in a two-tailed alpha of p = 0.025) that were larger than 97.25% of the random group assignments, were considered significantly different between genotypes.  This method avoids statistical assumptions about the data and corrects for multiple comparisons.

**Supplementary Figures:**

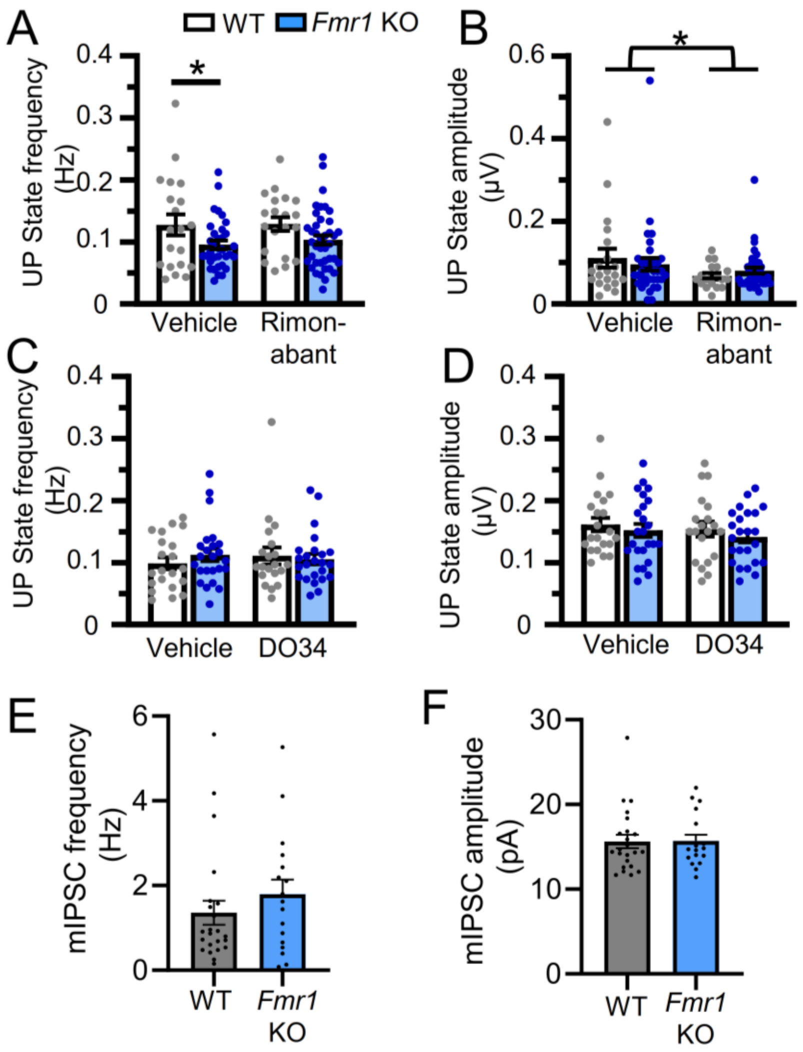


**Supplementary Figure 1. Effects of rimonabant and DO34 on frequency and amplitude of Up states and baseline mIPSC values.** A,B)  Rimonabant has no effect on Up state frequency (A), but generally reduces Up state amplitude (B) in WT and *Fmr1* KO slices.  2 way ANOVA; Main effect of rimonabant; F (1, 109) = 4.07; p< 0.05).  C,D) DO34 has no effect on Up state frequency or amplitude.  E,F) **Normal mIPSC frequency and amplitude in *Fmr1* KO L2/3 cortical neurons** E) Mean (±SEM) mIPSC frequency with values from individual L2/3 cortical neurons overlaid from WT and *Fmr1* KO mice F) Mean (±SEM) mIPSC amplitude with values from individual L2/3 cortical neurons overlaid from WT and *Fmr1* KO mice. (n= 23 WT; 17 *Fmr1* KO).  mIPSC frequency values were averaged during the 5 min pre-rimonabant washon baseline period from the experiments in Fig. 1G.

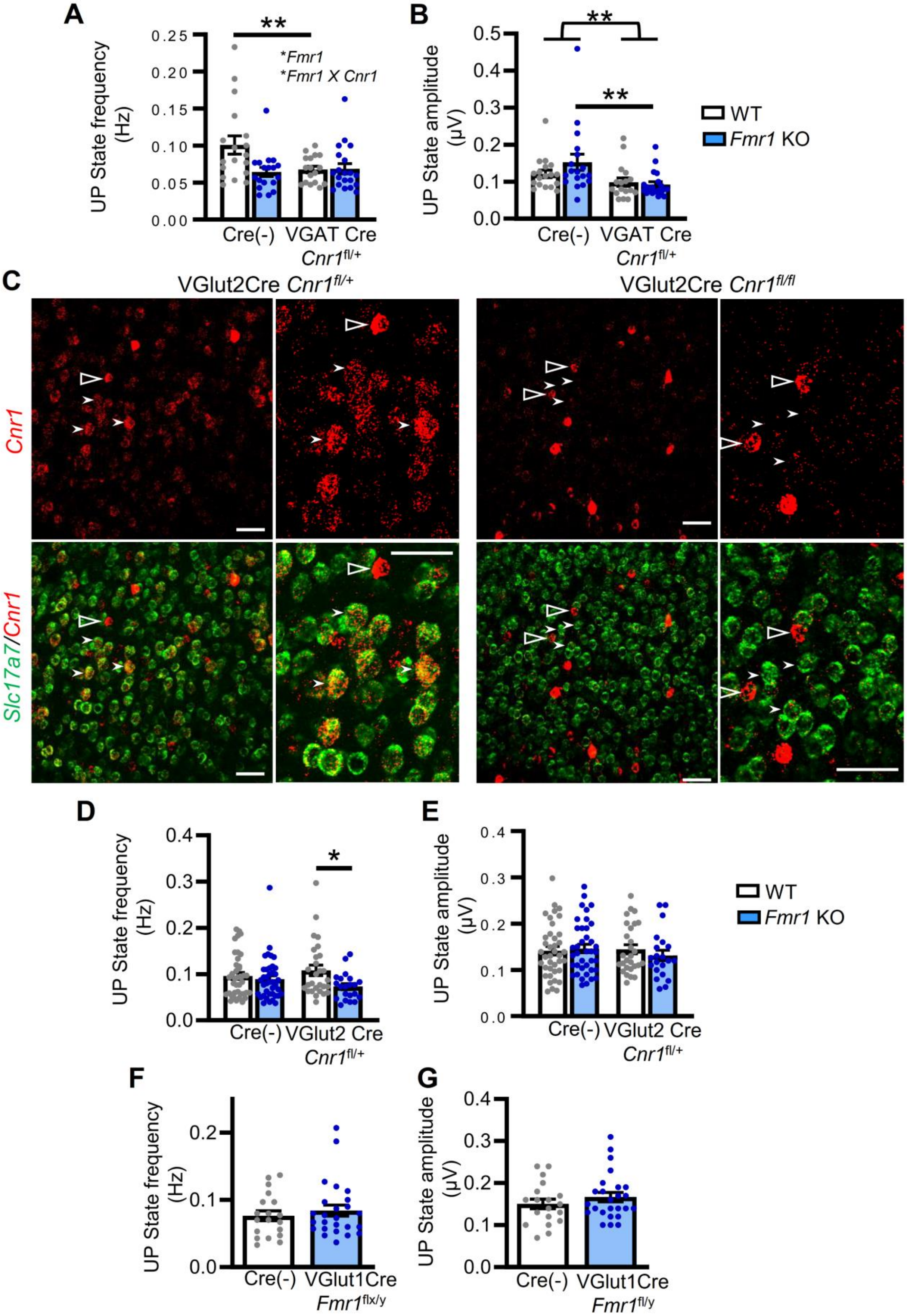


**Supplementary Figure 2. Effects of *Cnr1* and *Fmr1* deletion on Up state frequency and amplitude and**

**verification of *Cnr1* deletion in cortical glutamatergic neurons.** A,B) Heterozygous deletion of *Cnr1* in GABAergic neurons (VGATCre *Cnr1*$^{fl/+}$) reduced Up state frequency (A) in WT, but not *Fmr1* KO, slices and had a main effect on Up state amplitude; $F (1, 109) = 4.07$; $p< 0.05$). C) Reduction of *Cnr1* gene dosage in cortical glutamatergic neurons was accomplished by crossing the VGlut2-Cre and *Cnr1*$^{flx/WT}$ mouse lines. Images of homozygous deletion using RNAscope for *Cnr1* (Red) and the VGlut1 message, *Slc17a7*, (green) show our ability to effectively target gene dosage in glutamatergic neurons. While Cre-expression is specific for glutamatergic neurons in the VGlut2-Cre mouse, it is only expressed perinatally. Therefore, we used VGlut1 RNA, *Slc17a7*, for RNAscope to mark glutamatergic neurons. D,E) Heterozygous deletion of *Cnr1* in glutamatergic neurons (VGlut2Cre *Cnr1*$^{fl/+}$) did not affect Up state frequency (D) or amplitude (E). F,G) *Fmr1* deletion in glutamatergic neurons (VGlut1Cre *Fmr1* $^{fl/y}$) has no effect on Up state frequency (A) or amplitude (B). *, $p<0.05$ **, $p<0.01$. 2-way ANOVA with posthoc comparisons.

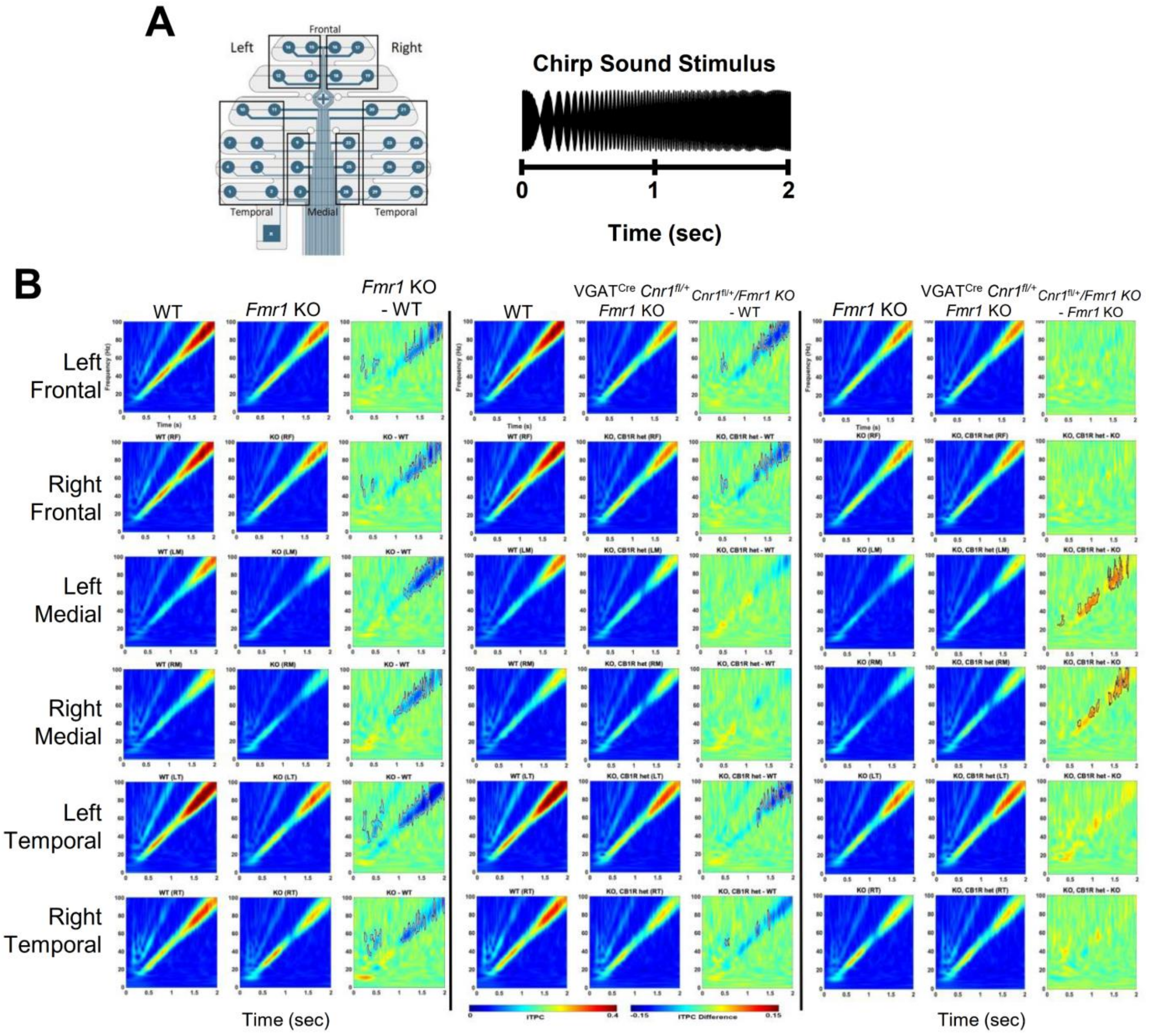


**Supplementary Figure 3. Genetic reduction of *Cnr1* in GABAergic neurons rescues decreased synchronization of chirp-driven cortical activity in medial recording locations as measured by EEG**. ITPC was quantified to measure the ability of sounds to consistently drive synchronous activity in cortex from trial to trial. A) *Left*: Diagram of EEG electrode array laid over scalp of mice with defined regions. *Right*: Chirp stimuli were 2 seconds of broadband noise sinusoidally modulated starting at 1 Hz and ramping up to 100 Hz. B) The data are divided into 3 vertical columns based on genotypic comparison (e.g. WT vs. KO on the left). Rows correspond to regions in the EEG electrode array, and ITPC plots are averages of individual electrodes in the region. In a single comparison column and single region row, two ITPC traces are on the left and a difference ITPC is plotted on the right. In the 2 medial regions, the difference plots most clearly indicate decreased ITPC at higher frequencies in *Fmr1* KO mice (left) and the rescue, or correction, of this phenotype in *Fmr1* KO with heterozygous deletion of *Cnr1* in GABAergic neurons (VGAT$^{Cre}$/*Cnr1*$^{fl/+}$/*Fmr1* KO). The ITPC deficit in the *Fmr1*

KO is not rescued in frontal or temporal regions. N=17,17,17 mice. Solid lines borders in the difference plots indicate statistically different areas of the plot.

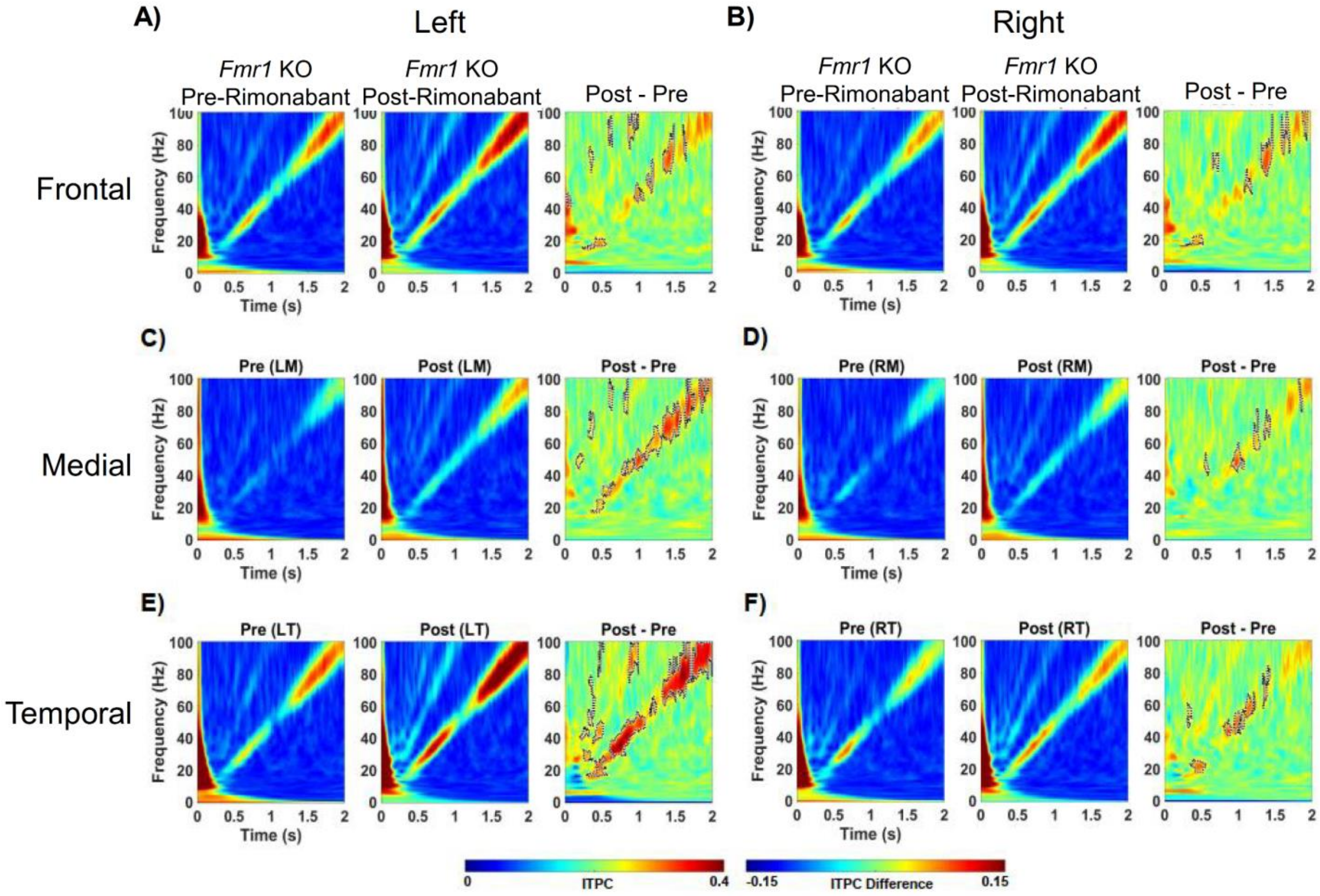


**Supplementary Figure 4. Rimonabant treatment (7 days) rescues decreased synchronization of chirp-driven cortical activity across brain regions as measured by EEG** A,B) ITPC average plots of induced activity for pre- and post-rimonabant treatment obtained from the Left (A) and Right (B) Frontal regions in *Fmr1* KO mice. *Far Right:* ITPC difference plot indicating increases in ITPC with rimonabant treatment. C,D) Same as in A and B, but for medial regions. E,F) Same as in A and B, but for temporal regions. N=14 mice. Solid lines borders in the difference plots indicate statistically different areas of the plot.

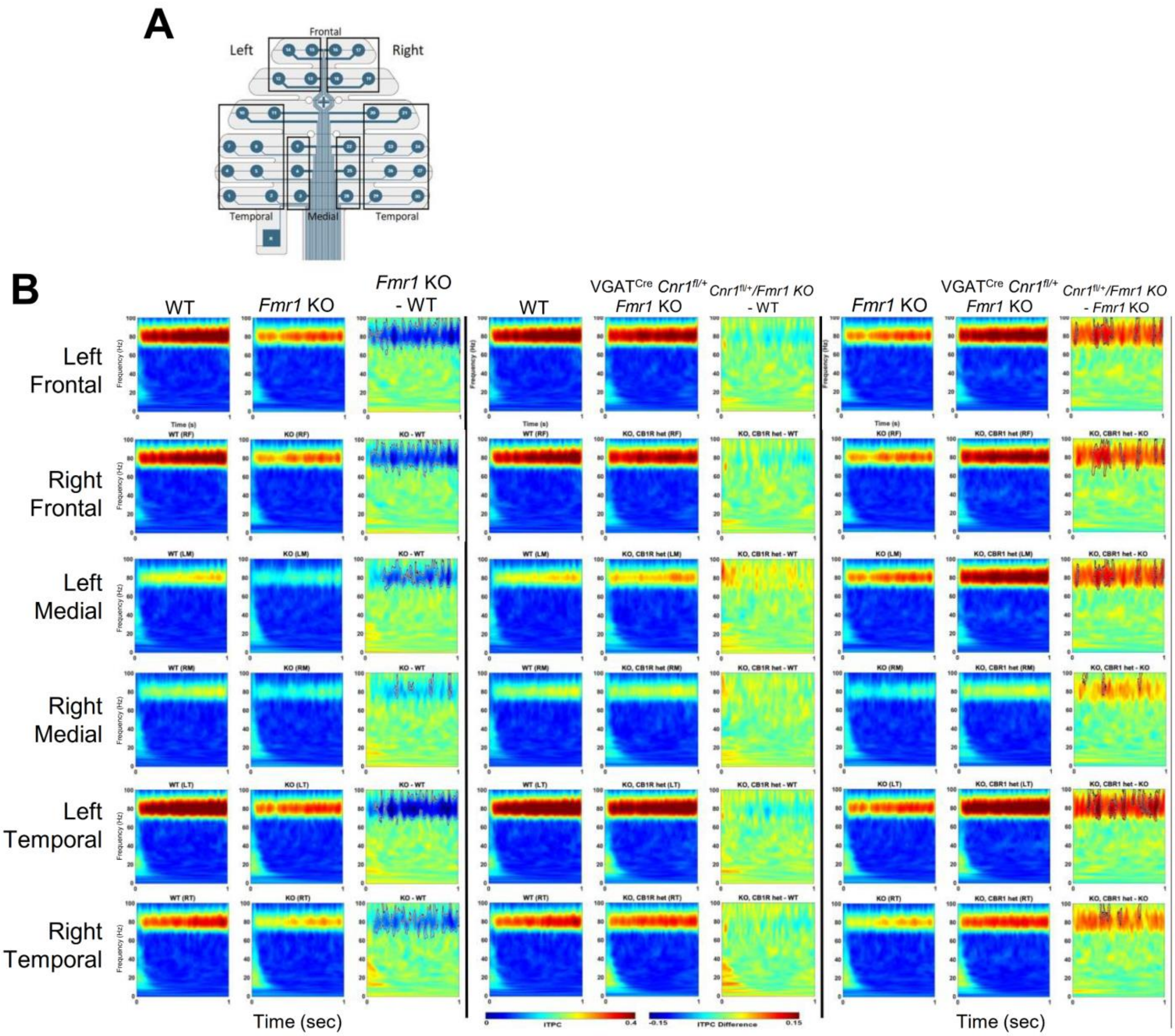


**Supplementary Figure 5. Genetic reduction of *Cnr1* in GABAergic neurons rescues decreased synchronization of pulse-driven cortical activity at 80 Hz (80 Hz ASSR) across all brain regions**. A) Diagram of EEG electrode array laid over scalp of mice with defined regions. B) An 80 Hz pulse train was used to induce an ASSR. In 5 out of 6 regions in the electrode array, difference plots indicate decreased ITPC of the 80 Hz ASSR in *Fmr1* KO mice and the rescue of this phenotype with heterozygous deletion of *Cnr1* in GABAergic neurons (VGAT$^{Cre}$/*Cnr1*$^{fl/+}$/*Fmr1* KO). N=17,17,17 mice. Solid lines borders in the difference plots indicate statistically different areas of the plot.

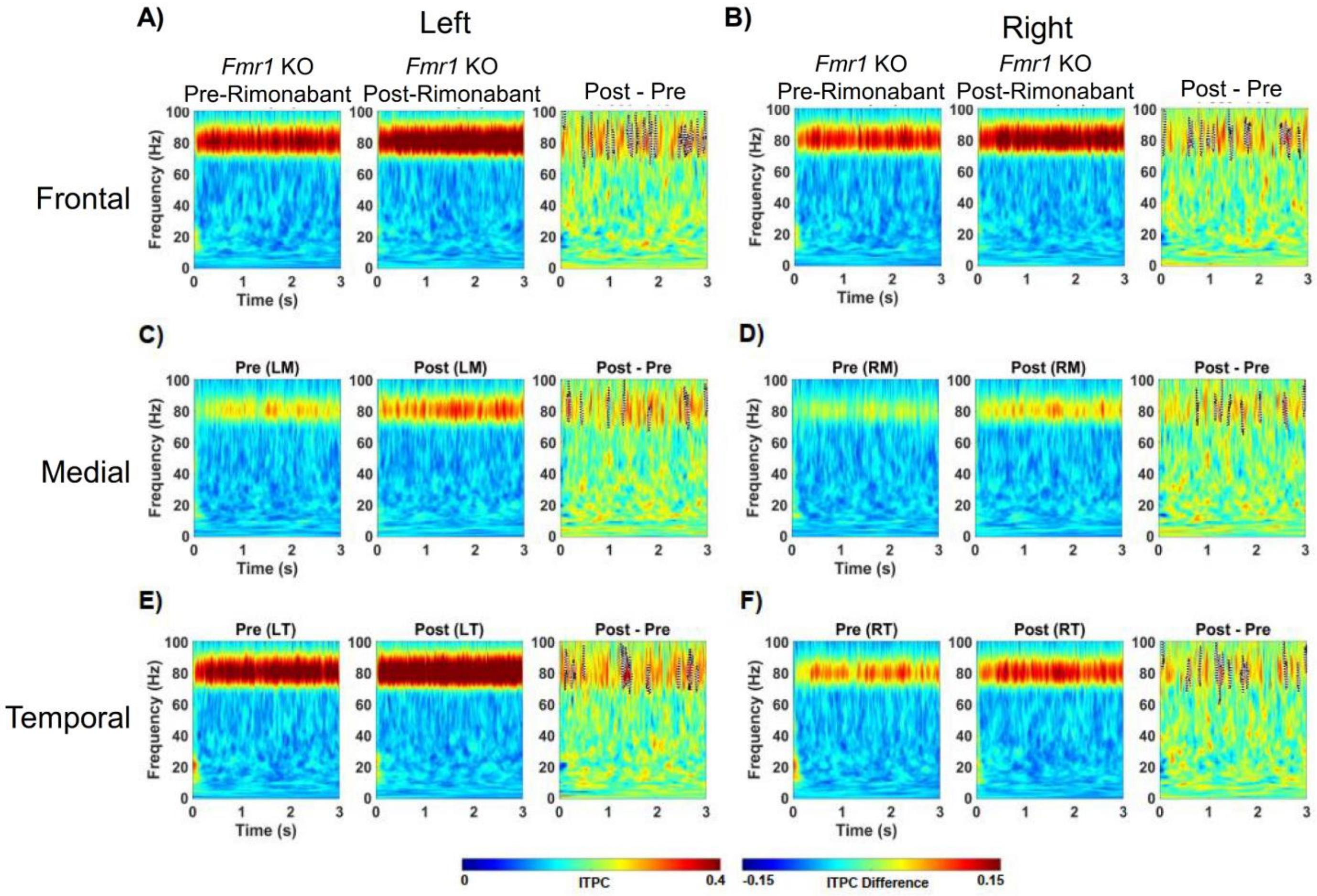


**Supplementary Figure 6. Rimonabant treatment (7 days) rescues decreased synchronization of pulse-driven cortical activity at 80 Hz (80 Hz ASSR)**. An 80 Hz pulse train was used to induce an ASSR. A,B) ITPC average plots of ASSR for pre- and post-rimonabant treatment obtained from the Left (A) and Right (B) Frontal regions in *Fmr1* KO mice. *Far Right:* ITPC difference plot indicating increases in ITPC with rimonabant treatment across all brain regions. C,D) Same as in A and B, but for medial regions. E,F) Same as in A and B, but for temporal regions. N=14 mice. Solid lines borders in the difference plots indicate statistically different areas of the plot.

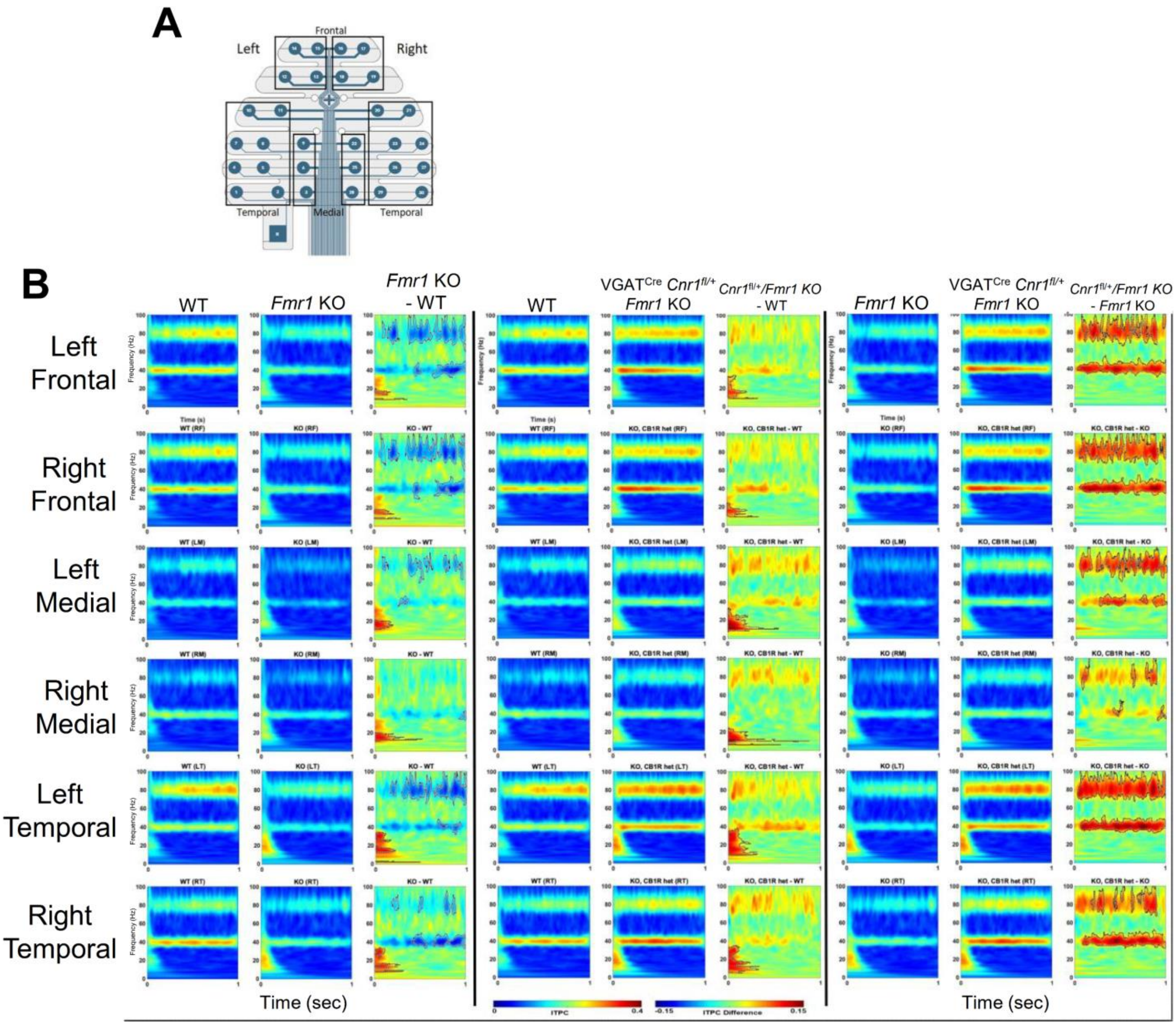


**Supplementary Figure 7. Genetic reduction of *Cnr1* in GABAergic neurons rescues decreased synchronization of pulse-driven cortical activity at 40 Hz (40 Hz ASSR) across all brain regions**. A) Diagram of EEG electrode array laid over scalp of mice with defined regions. B) A 40 Hz pulse train was used to induce an ASSR. In 5 out of 6 regions in the electrode array, difference plots indicate decreased ITPC of the 40 Hz ASSR in *Fmr1* KO mice and the rescue of this phenotype in *Fmr1* KO mice with heterozygous deletion of *Cnr1* in GABAergic neurons (VGAT$^{Cre}$/*Cnr1*$^{fl/+}$/*Fmr1* KO). N=17,17,17 mice. Solid lines borders in the difference plots indicate statistically different areas of the plot.

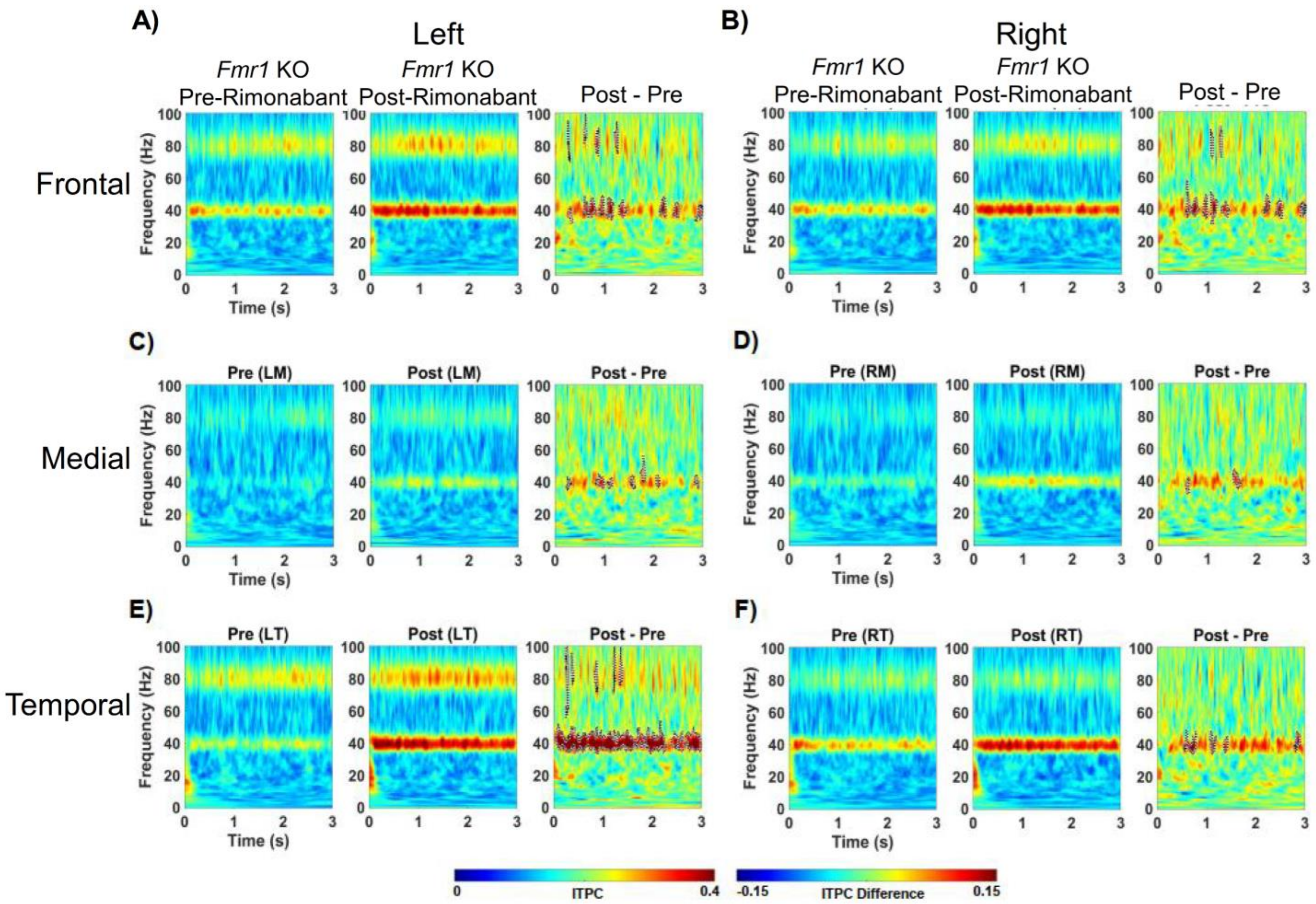


**Supplementary Figure 8. Rimonabant treatment (7 days) rescues decreased synchronization of pulse-driven cortical activity at 40 Hz (40 Hz ASSR)**. A 40 Hz pulse train was used to induce an ASSR. A,B) ITPC average plots of ASSR for pre- and post-rimonabant treatment obtained from the Left (A) and Right (B) Frontal regions in *Fmr1* KO mice. *Far Right:* ITPC difference plot indicating increases in ITPC with rimonabant treatment across all brain regions. C,D) Same as in A and B, but for medial regions. E,F) Same as in A and B, but for temporal regions. N=14 mice. Solid lines borders in the difference plots indicate statistically different areas of the plot.

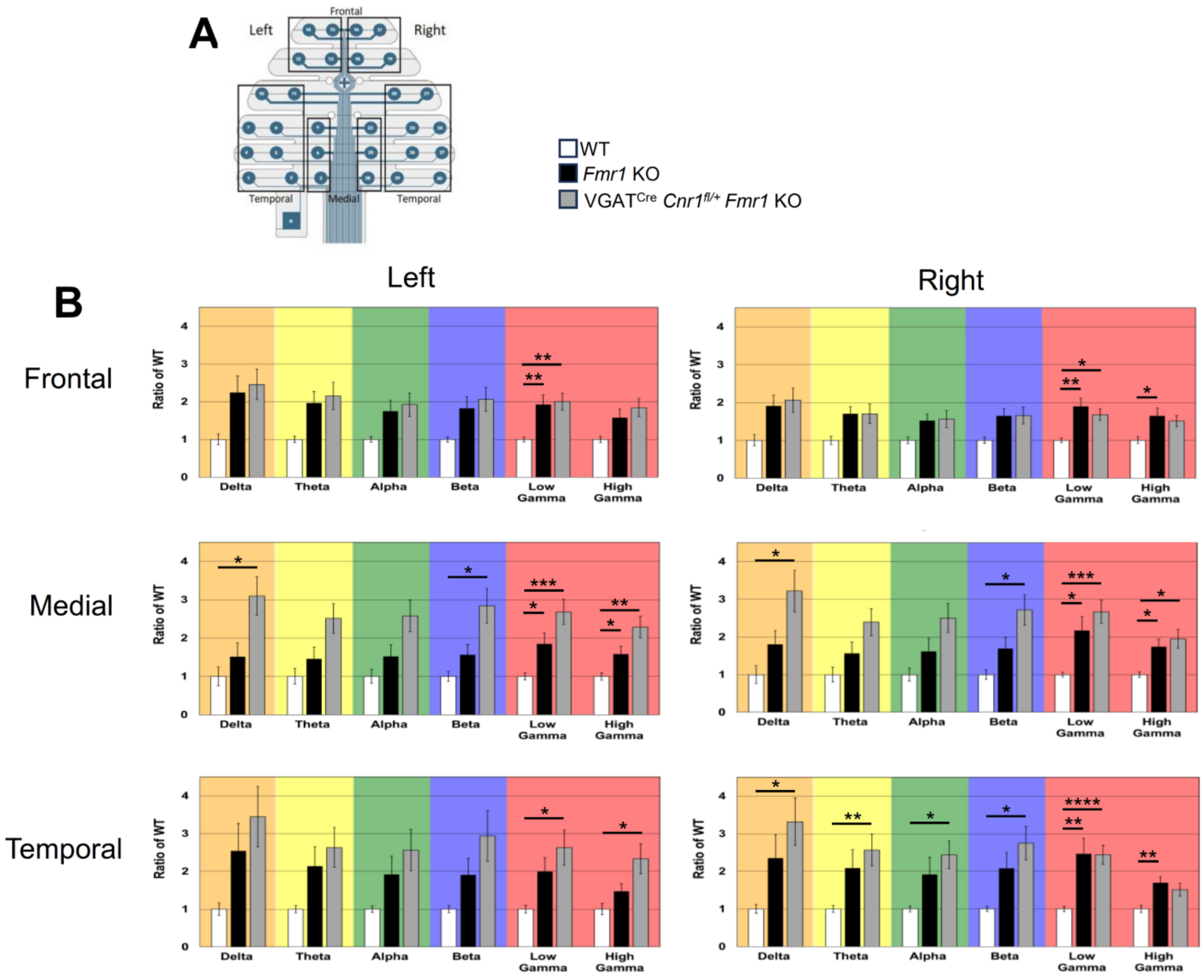


**Supplementary Figure 9. Genetic reduction of *Cnr1* in GABAergic neurons does not rescue increased resting state gamma power in the *Fmr1* KO mouse *in vivo***. Power spectral density normalized to wild-type mice is plotted for 6 standard frequency bands. Increases in power are observed across in the "low gamma" frequency band (30-55Hz) across many brain regions in the *Fmr1* KO, and this increase is not rescued in *Fmr1* KO with heterozygous deletion of *Cnr1* in GABAergic neurons (VGAT$^{Cre}$/*Cnr1*$^{fl/+}$/*Fmr1* KO). There were no differences between *Fmr1* KO and VGAT$^{Cre}$/*Cnr1*$^{fl/+}$/*Fmr1* KO mice at any frequencies in any brain region. N=17, 17, 17 mice. * p<0.05, ** p<0.01 , *** p<0.001, ****p< 0.0001.

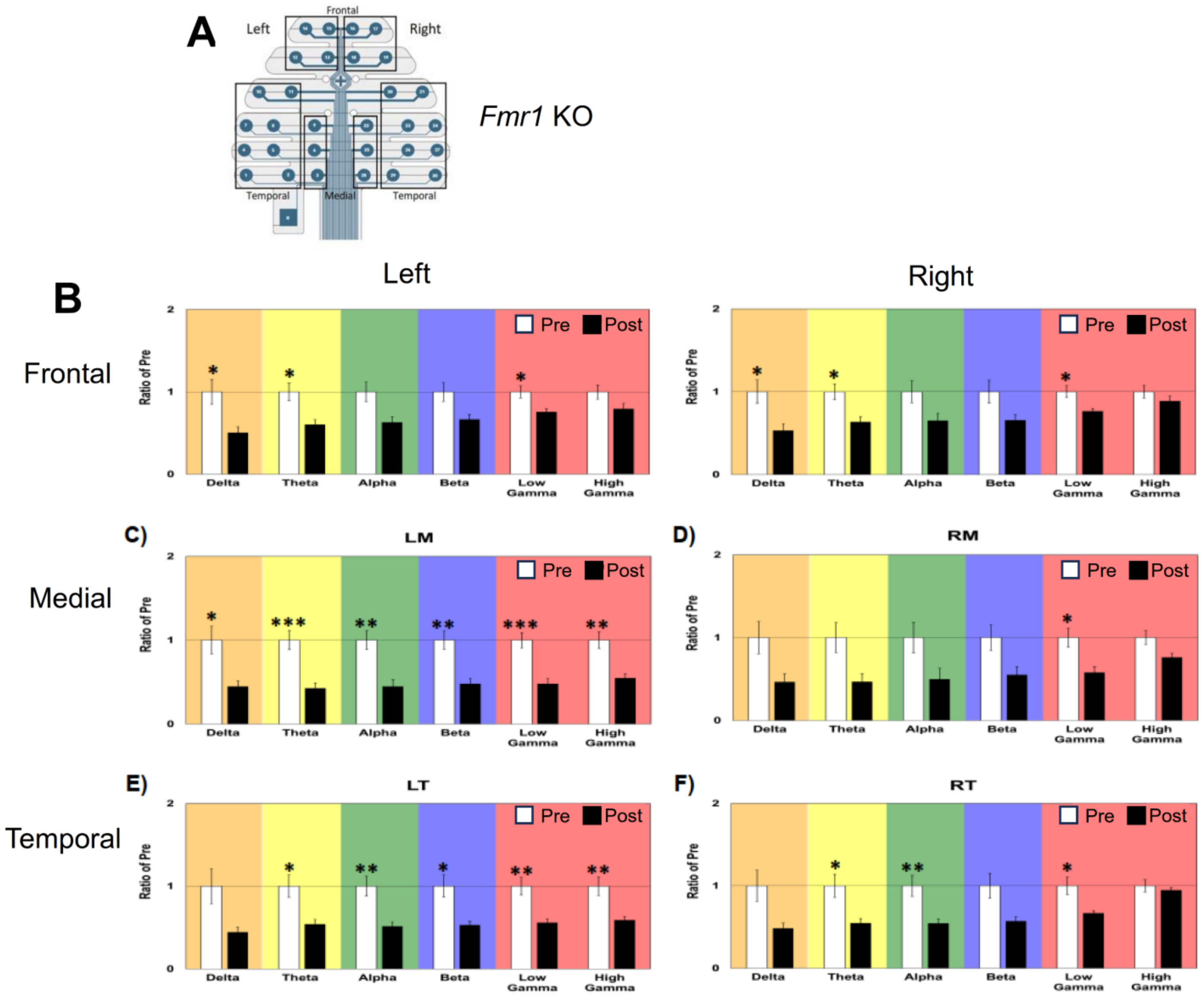


**Supplementary Figure 10. Rimonabant treatment (7 days) reduces resting power across many frequency bands in the *Fmr1* KO mouse *in vivo*.** Power in 6 standard frequency bands is normalized to *Fmr1* KO mice before rimonabant treatment (pre). 7 day rimonabant treatment of *Fmr1* KO mice (post) decreases power in many frequency bands across multiple brain regions. N=14 mice. * p<0.05, ** p<0.01 , *** p<0.001.
